# Basal forebrain gating by somatostatin neurons drives cortical activity

**DOI:** 10.1101/102582

**Authors:** Nelson Espinosa, Alejandra Alonso, Cristian Morales, Pablo Fuentealba

## Abstract

The basal forebrain provides modulatory input to the cortex regulating brain states and cognitive processing. Somatostatin-expressing cells constitute a local GABAergic source known to functionally inhibit the major cortically-projecting cell types. However, it remains unclear if somatostatin cells can regulate the basal forebrain’s synaptic output and thus control cortical dynamics. Here, we demonstrate in mice that somatostatin neurons regulate the corticopetal synaptic output of the basal forebrain impinging on cortical activity and behavior. Optogenetic inactivation of somatostatin neurons in vivo increased spiking of some basal forebrain cells, rapidly enhancing and desynchronizing neural activity in the prefrontal cortex, inhibiting slow rhythms and increasing gamma oscillations. Locomotor activity was specifically increased in quiescent animals, but not in active mice. Altogether, we provide physiological and behavioral evidence indicating that somatostatin cells are pivotal in gating the synaptic output of the basal forebrain, thus indirectly controlling cortical operations via both cholinergic and non-cholinergic mechanisms.

## Introduction

The mammalian basal forebrain is a collection of subcortical structures comprising the ventral pallidum, diagonal band of Broca, substantia innominata, medial septum and peripallidal region, which provides extensive axonal projections to the entire cerebral cortex (Jones, 2008, Zaborszky et al., 2012). Damage to the basal forebrain is of conspicuous relevance for several neurological disorders, including Alzheimer’s disease, Parkinson’s disease, schizophrenia, and drug abuse (Whitehouse et al., 1982, Conner et al., 2003, Smith et al., 2004). Under normal physiological conditions, the basal forebrain plays central roles in arousal, attention, motivation, memory, plasticity, sensory processing and sleep-wake cycles (Pinto et al., 2013, Lin et al., 2015, Xu et al., 2015). These actions are achieved by the complementary roles of a heterogeneous mixture of cell types that differ in neurotransmitter content, somato-dendritic morphology, axonal projections and spike timing (Brashear et al., 1986, Zaborszky and Duque, 2000, Jones, 2005).

Despite representing a minor fraction of the basal forebrain neuronal population, cholinergic projection cells have been extensively studied and implicated in most of the abovementioned functions. Nonetheless, in recent years evidence has emerged on the functional significance of different non-cholinergic cells in the basal forebrain, which include neuronal populations expressing GABA, glutamate and neuropeptides (Duque et al., 2000, Zaborszky and Duque, 2000, Henny and Jones, 2008). Recent comprehensive circuit-mapping experiments have established the hierarchical organization of the basic synaptic circuit of sleep-wake cycle in the basal forebrain. Accordingly, the main three cortically projecting cell types are synaptically connected, with glutamatergic cells exciting cholinergic neurons, which in turn activate parvalbumin-expressing cells (Xu et al., 2015). The activation of this circuit exerts a prominent wake-promoting effect by desynchronization of cortical activity, a hallmark of wakeful and alert brain states enhancing cortical responsiveness and sensory encoding (Goard and Dan, 2009, Pinto et al., 2013, Xu et al., 2015). In particular, non-cholinergic glutamatergic neurons showed the strongest wake-promoting effect, consistent with their hierarchical position in the circuit. Conversely, optogenetic activation of basal forebrain somatostatin-expressing neurons rapidly increased the probability of slow wave sleep, with several of these neurons being strongly active during that brain state (Xu et al., 2015). Furthermore, somatostatin neurons in the basal forebrain actively inhibit all three major types of wake-promoting neurons. Thus, promotion of slow wave sleep seems to be based on the broad inhibition of multiple wake-promoting cell types in the basal forebrain local circuit.

Optogenetic stimulation of somatostatin neurons in the basal forebrain with Channelrhodopsin-2 has demonstrated that they are *sufficient* to promote deep sleep (Xu et al., 2015). However, it remains unknown if they are *necessary* to regulate cortical dynamics. Here, we address this issue through optogenetic inactivation of somatostatin neurons. We found that somatostatin neurons could control the corticopetal synaptic output of the basal forebrain affecting the intrinsic dynamics of cortical circuits in anesthetized and freely-moving mice. Selective inhibition of somatostatin neurons rapidly increased neural activity in a subset of basal forebrain cells, followed by enhanced recruitment of cortical cells and desynchronization of prefrontal cortex activity. These results suggest that somatostatin neurons are a key element in the control of the synaptic output of the basal forebrain and can thus affect the regulation of cortical states.

## Results

### Synaptic output of the basal forebrain is regulated by somatostatin cells

We first studied the significance of somatostatin cells for basal forebrain activity patterns. For this, we stereotaxically implanted an optrode in the basal forebrain of anesthetized transgenic animals (Fig. 1A) selectively expressing halorhodopsin in somatostatin cells (NpHR+, supplementary figure 1). We delivered prolongued laser pulses to achieve maximal inhibition of somatostatin cells and reproduce previous experimental protocols (Goard and Dan, 2009, Pinto et al., 2013, Kim et al., 2016). We found a minor fraction of basal forebrain cells (8.8%, n = 29 units) responding (time constant = 263.1 ± 48.7 ms) by robustly decreasing their activity (49.2 ± 5.1%). Along with these putative somatostatin cells decreasing their firing rate, another neuronal population (17%, n = 56 units) increased its activity (39.6 ± 7.9 %), presumably by synaptic disinhibition, with significantly slower kinetics (time constant = 602.5 ± 68.6 ms) (Fig. 1C, E). Spontaneous firing rates of neurons activated by optical stimulation were consistently higher than those of inhibited or unresponsive cells (Fig. 1D), suggesting that they might belong to different cell classes (Lee et al., 2005, Hassani et al., 2009). Thus, optical inactivation of somatostatin neurons was rapidly followed by the activation of a subset of basal forebrain cells.

**Figure 1.**
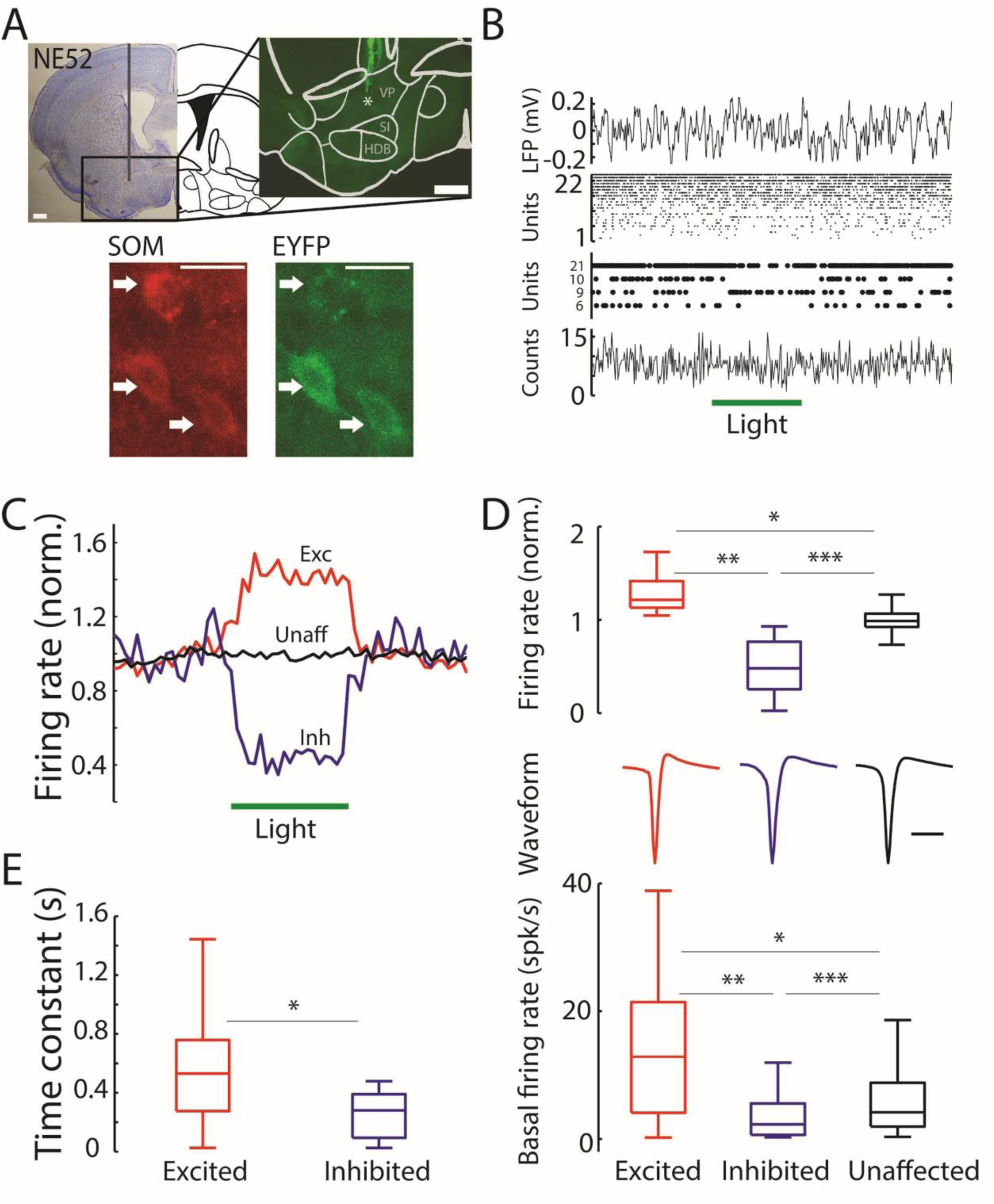
Optogenetic inactivation of somatostatin cells in the basal forebrain. **A**, Nissl stained brain section and schematic coronal drawing of the mouse brain (mouse NE52, section 46, adapted from (Franklin and Paxinos, 2007)). Scale bar: 400 um. Inset: fluorescence microscopy photomicrograph of basal forebrain region with optrode stained with DiI (arrow). Gray lines depict anatomical nuclei. Scale bar: 200 um. Bottom, fluorescence microscopy photomicrograph shows a cluster of cells co-expressing NpHR (EYFP) and somatostatin (SOM) in the basal forebrain. **B**, electrophysiological recordings from the brain location shown in (A). From top to bottom: LFP (filtered 0.1 Hz - 5 kHz), raster plots for all simultaneously recorded cells, raster plot for laser-responsive units, and multiunit histogram. Binsize: 5 ms. **C**, average normalized discharge probability for excited (Exc., red line, n = 56), inhibited (Inh., blue line, n = 29), and unaffected (Unaff., black line, n = 245) neurons recorded in the basal forebrain (n = 4 animals). Horizontal bar depicts laser stimulation (5 s, 4-6 mW fiber diameter 100 um). Binsize: 250 ms. **D**, normalized discharge probability during optical stimulation (top panel, Kruskal-Wallis test, *p* = 5.44×10^-35^; Wilcoxon rank-sum test, \**p* = 5.65×10^-24^, \*\**p* = 5.39×10^-14^, \*\*\**p* = 1.11×10^-16^), spike waveform average (middle panel), and basal firing rate (bottom panel, Kruskal-Wallis test, *p* = 2.06×10^-8^; Wilcoxon rank-sum test, \**p* = 8.17×10^-8^, \*\**p* = 4.10×10^-5^,\*\*\**p* = 0.0125) for excited, inhibited, and unaffected cells; respectively. Scale bar: 1 ms. **E**, exponential time constants from curve fittings to the early response of neurons excited or inhibited by optical stimulation (Wilcoxon rank-sum test, \**p* = 0.0072).

We then combined somatosensory stimulation with optical inhibition of somatostatin cells in order to physiologically characterize response patterns in the basal forebrain. We found neuronal patterns consistent with different cell types being engaged by optical stimulation (supplementary figure 2). Previous studies have shown that only a subset of non-cholinergic cells is inhibited during somatosensory stimulation by tail pinching (Hassani et al., 2009). Similarly, we found a group of cells inactivated by somatosensory stimulation, which exhibited diverse responses to optical stimulation, suggesting functional diversity among them. Indeed, putative non-cholinergic cells were excited, inhibited or unaffected by optical stimulation. It has also been documented that cholinergic cells display low levels of activity during slow oscillations and are strongly activated during somatosensory stimulation (Lee et al., 2005, Hassani et al., 2009). We found units consistent with such activity patterns that were also disinhibited by the inactivation of somatostatin cells, thus suggesting that some cholinergic cells were also recruited by optical stimulation. Finally, somatostatin cells exhibited diverse response patterns to somatosensory stimulation, consistent with the diverse firing patterns described across the sleep-wake cycle (Xu et al., 2015). Overall, our results suggest that decreasing the GABAergic input provided by somatostatin neurons in the basal forebrain modifies the balance in network activity engaging diverse neuronal populations, likely comprising cholinergic and non-cholinergic cortically projecting cells.

**Figure 2.**
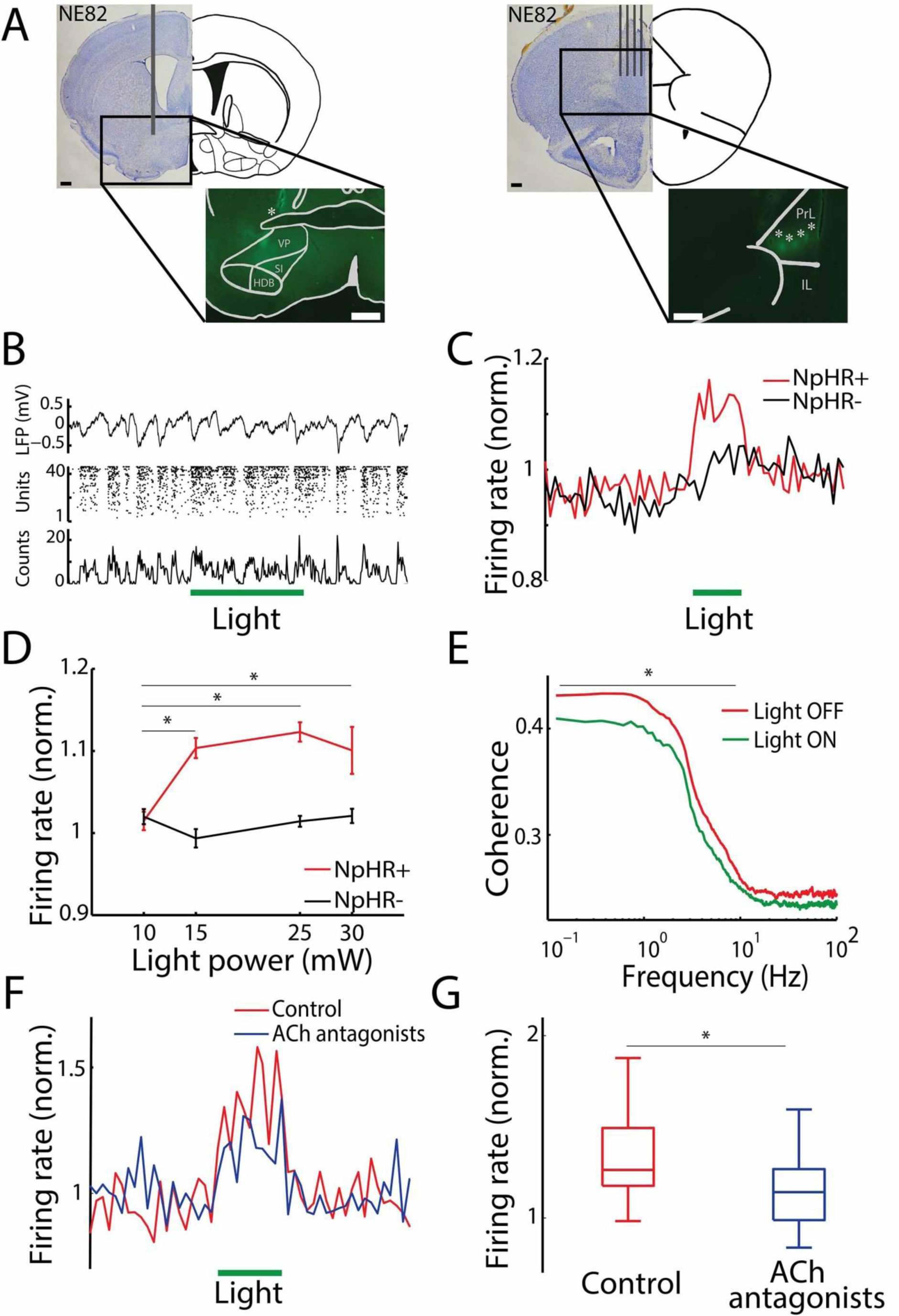
Neuronal activity in the prefrontal cortex during optical inhibition of basal forebrain somatostatin cells. **A**, photographic montage of Nissl stained brain sections and schematic coronal drawing. Left, optical fiber location in the basal forebrain (gray vertical line) (mouse NE82; left, section 46, adapted from (Franklin and Paxinos, 2007)). Scale bar: 400 um. Inset: fluorescence microscopy photomicrograph of basal forebrain region with optical fiber tract stained with DiI. Gray lines depict nuclei on atlas section. Scale bar: 200 um. Right: silicon probe location (brain NE82; right, section 16, adapted from (Franklin and Paxinos, 2007)). Scale bar: 400 um. Inset: fluorescence microscopy of prefrontal cortex with silicon probe stained with DiI. Gray lines depict prefrontal cortex borders on atlas section. Scale bar: 200 um. **B**, example electrophysiological recording from the brain shown in (A). Top panel: LFP; middle panel: raster plot for all recorded units; bottom panel: multiunit histogram. Binsize: 4 ms. **C**, normalized discharge probability averages for animals expressing functional halorhodopsin (NpHR+, red line, n = 1308 units, 7 animals) and control animals (NpHR-, black line, n = 851 units, 3 animals). Optogenetic stimulation produced significantly different responses (*W* = 74742, *p* < 10^-6^, Wilcoxon signed-rank test). Horizontal bar indicates optical stimulation (5 s, 15-25 mW). Binsize: 500 ms. Note slower and smaller response in control animals (NpHR-), likely due to temperature effects (supplementary figure 4). Note nonspecific neuronal activation in NpHR- animals, which was likely due tissue heating from laser stimulation (Stujenske et al., 2015). **D**, Plot depicting average discharge probability versus light power (NpHR+, n = 9 animals; NpHR-, n = 5 animals). Error bars, s.e.m. *p < 0.001 (Two-way ANOVA test and Bonferroni *post hoc* correction). **E**, Average coherence between single unit and multi-unit activity in the presence (green line, light on) or absence (black line, light off) of optogenetic stimulation. Laser-induced reduction of coherence was statistically significant only for low frequencies (< 10 Hz, *W* = 142155, *p* < 10^-6^, Wilcoxon signed rank test, n = 1109 cells, 7 NpHR+ animals). **F**, average normalized discharge probability of cortical neurons before (black line) and after (blue line) the local injection of cholinergic blockers (200 nl, 2 mM, atropine and mecamylamine) in the prefrontal cortex Binsize: 500 ms. Horizontal green line depicts laser stimulation (5 s, 15 - 25 mW). **G**, normalized discharge probability during optical stimulation before (black box) and after (blue box) the local injection of cholinergic blockers (W = 1438, *p* < 10^-6^, Wilcoxon signed rank test, n = 64 cells, 3 animals).

### Optogenetic disinhibition of the basal forebrain drives cortical dynamics

Next, we assessed the effect of the basal forebrain’s synaptic output onto cortical neurons. Hence, we implanted an optic fiber in the basal forebrain and simultaneously recorded neural activity in the medial prefrontal cortex (mPFC, Fig. 2A). Optogenetic inhibition of somatostatin cells in the basal forebrain reliably increased mPFC spiking activity (Fig. 2B, C). Nearly one third of cortical neurons (29.6%, n = 387) systematically increased their firing rate (by 30.4 ± 1.2%) for the entire duration of the laser pulse, producing a prominent effect on cortical activity (Fig. 2C). Enhanced discharge probability was slightly higher in infralimbic cortex as compared with prelimbic cortex (supplementary figure 3), consistent with differential connectivity patterns provided by corticopetal basal forebrain projections (Henny and Jones, 2008). The excitatory effect was dependent on laser power, and specific for transgenic NpHR+ animals (Fig. 2C, D; supplementary figure 4). Moreover, optogenetic stimulation of transgenic NpHR+ animals was only effective in evoking cortical activation, when the optic fibre was accurately positioned in the basal forebrain, and not in other brain regions (supplementary figure 5). Thus, the effect of cortical activation was specific for optogenetic disinhibition of the basal forebrain. Laser induced cortical activation also caused a marked reduction of neural synchrony measured by the coherence between individual neurons (i.e., single units) and the other simultaneously recorded cells (i.e., multiunits), in particular for activity in the low frequency range (< 10 Hz, Fig. 2E). Previous studies have shown that enhanced and decorrelated cortical activity can attributed to the activation of basal forebrain cholinergic pathways innervating the cortex (Goard and Dan, 2009, Pinto et al., 2013). In order to confirm that cholinergic projection neurons mediated laser induced cortical activation in our experimental conditions, we locally applied cholinergic receptor antagonists in the medial prefrontal cortex. Cholinergic blockers significantly diminished cortical activation (from 35.7 ± 3.2 % (before drug), to 19.0 ± 3.7 % (after drug); W = 1438, *p* < 10^-6^, Wilcoxon signed rank test, n = 64 cells, 3 mice), confirming that at least part of the effect of basal forebrain activation was mediated by enhanced cholinergic transmission to the cortex. Importantly, baseline spiking activity in the cortex was not affected by cholinergic receptor antagonists. On the other hand, optical stimulation produced a significant increase in discharge probability at all intervals (Friedman test, p = 1.07×10^-18^). Nevertheless, the effect of optical stimulation progressively decreased after local drug application (Wilcoxon signed rank test, p = 2.6 × 10^-4^). In order to confirm that cholinergic projection neurons mediated laser induced cortical activation, we locally applied cholinergic receptor antagonists in the medial prefrontal cortex. Cholinergic blockers significantly diminished cortical activation (from 35.7 ± 3.2%, before drug; to 19.0 ± 3.7%, after drug), confirming that at least part of the effect of basal forebrain activation was mediated by enhanced cholinergic transmission to the cortex (supplementary figure 5).

**Figure 3.**
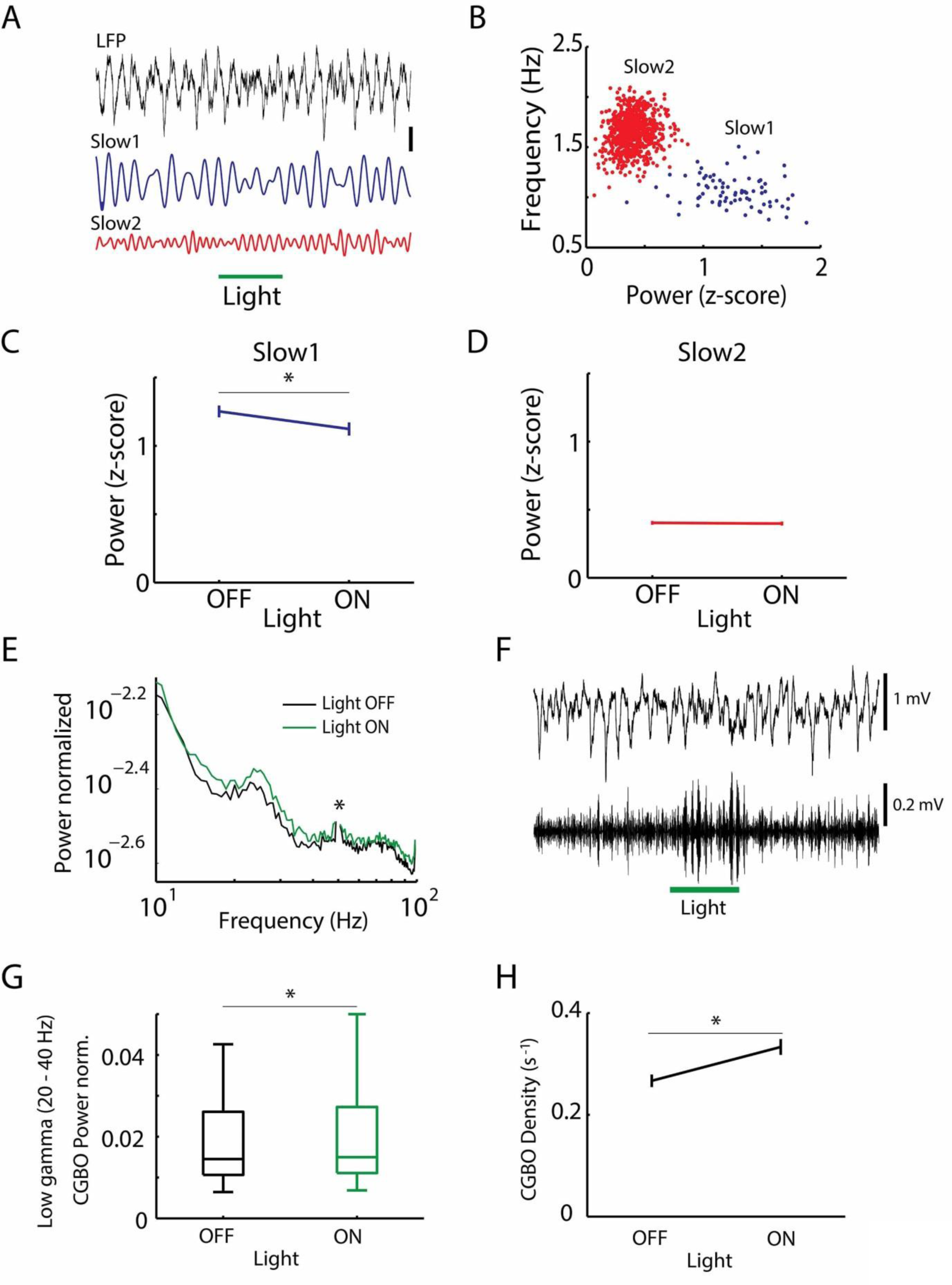
Cortical oscillations during optical inhibition of basal forebrain somatostatin cells. **A**, example of LFP (filtered 0.1 Hz – 5 kHz, top panel) and slow oscillations in two frequency bands (slow 1, filtered 0.5-1 Hz, middle panel; and slow 2, bottom panel, filtered 1-2 Hz) from cortical activity. Horizontal green line denotes laser stimulation (5 s, 25 mW). Scale bar: 0.4 mV. Note that only slow 1 oscillations are affected by light. **B**, scatter plot of the normalized power versus main frequency of slow oscillations (n = 870 epochs, 7 animals). Two frequency bands were identified by cluster analysis (see Methods), i.e., slow 1 (blue dots) and slow 2 (red dots). **C**, normalized power for slow 1 oscillations in the presence (on) and absence (off) of optical stimulation. Gray lines indicate individual values (n = 77). Black line represents mean ± s.e.m (paired t-test, \**p* = 0.0009316). **D**, normalized power for slow 2 oscillations in the presence (on) and absence (off) of optical stimulation (n = 793, mean ± s.e.m, paired t-test, *p* = 0.22603). **E**, average power spectral distribution of cortical LFP (n = 7 NpHR+ animals) in the presence (green line) or absence (black line) of laser stimulation. Spectral distribution was normalized by 1/f factor (see Methods). Asterisk (*) depicts 50 Hz-artifact removal. **F**, example of wideband (filtered 0.1 Hz – 5 kHz, top panel) and gamma band (filtered 20 – 80 Hz, bottom panel) cortical LFP (NE49 reg01 pulse9 chan19). Horizontal green line shows laser stimulation (5 s, 25 mW). **G**, box plots for the average power of low gamma oscillations (20–40 Hz, \**p* = 0.0018, Wilcoxon signed-rank test). For high gamma oscillations (55-80 Hz) see supplementary figure 8. **H**, Plots (mean ± s.e.m.) show the density of low gamma episodes (paired t-test, \**p* = 5.76×10^-8^).

The multielectrode probe used in our experiments allowed us to simultaneously sample neural activity from different cortical layers (supplementary figure 6). We found that superficial layers contained a tiny fraction of the recorded cells, with the large majority of active cells (79.8 %) being located in deeper layers. Interestingly, despite large differences in the total numbers of cells by layer, similar proportions were activated at all depths. We next tested if optical stimulation of the basal forebrain exerted differential effects according to cortical cell type (supplementary figure 6). For this, we sorted units according to spike duration. A histogram of spike durations for all recorded units showed a bimodal distribution, with fast-spiking cells (i.e., spike duration < 0.6 ms), putative GABAergic interneurons (McCormick et al., 1985), accounting for a small fraction of the total neuronal population (9.9 %). Fast-spiking cells discharged at significantly higher rates (unpaired t-test, *p* = 3.9 × 10^-36^, 5.16 ± 0.39 Hz) than regular-spiking cells, (i.e., spike duration > 0.6 ms, 2.5 ± 0.05 Hz), putative pyramidal cells (McCormick et al., 1985). Optical stimulation of the basal forebrain activated similar proportions of putative interneurons and pyramidal cells. However, the increase in discharge probability produced by laser stimulation was significantly higher on interneurons (35.5 ± 5.9 %) than on pyramidal cells (26.2 ± 1.0 %; unpaired t-test, p = 0.019). Hence, the cell type-specific increase in discharge probability produced by laser stimulation suggests differential dynamics of neuronal activation by cell type in the cortex upon basal forebrain excitatory drive.

### Optogenetic stimulation of the basal forebrain reorganizes cortical oscillatory patterns

We sought to establish if the prominent effect in cortical spiking during basal forebrain stimulation was correlated with specific changes in network activity patterns in the cortex. Our recordings were performed in the context of anesthesia-induced slow oscillations, which resemble slow waves occurring during natural deep sleep and powerfully phase-modulate neuronal activity across the entire cortical mantle (Steriade et al., 1993a, Steriade, 2006). We normalized the LFP signal and plotted the distribution of baseline slow oscillation epochs in reference to laser stimulation (Fig. 3A, B). Hence, we found that slow oscillatory episodes distributed in two clusters (Fig. 3B), from which, only one cluster was affected by optical stimulation in the basal forebrain (Fig. 3C, D). That is, only very slow frequency, high power oscillations were suppressed by optogenetic stimulation. We then studied if slow oscillatory cortical activity was able to bias the effect of the input provided by the basal forebrain. We found that the effect of optogenetic stimulation on cortical activity was phase-modulated by slow oscillations (supplementary figure 7). Although basal forebrain disinhibition was able to enhance cortical activity in all phases of the slow oscillation, the effect was maximal during the active phase of slow oscillations and minimal at the peak of slow oscillation cycles, corresponding to the silent phase of the rhythm (supplementary figure 7). Furthermore, the influence of basal forebrain on cortical spiking did not extend for the whole duration of the light pulse (supplementary figure 7). Instead, optogenetic stimulation of the basal forebrain was phase modulated by the slow oscillation only during the second half of the laser pulse (i.e., 3-5 s, supplementary figure 7). These results suggest that the impact of basal forebrain output on cortical activity strongly depends on the ongoing cortical state.

In addition, we analyzed cortical gamma band activity, a prominent marker of cortical activation (Sohal et al., 2009), which seems to rely on the activity of basal forebrain parvalbumin-expressing cells, rather than depend on cholinergic neurons (Kim et al., 2015). Hence, we quantified the spectral distribution of cortical activity and found a prominent shoulder in the low gamma range (20-40 Hz, Fig. 3E), which power was significantly more elevated during optical stimulation when compared to control periods (Wilcoxon signed rank test, p = 0.0226, Fig. 3G). This result was specific for the low gamma band, as it was not detected in the high gamma band (55-80 Hz, supplementary figure 8). In addition, we also computed the density of gamma oscillatory episodes (Sirota et al., 2008, Le Van Quyen et al., 2010, Valderrama et al., 2012). Accordingly, we analytically extracted gamma band episodes from the gamma band activity. We found that the density of oscillatory episodes significantly increased (t-test, p = 5.8 × 10^-8^) only in the low gamma band during optical stimulation (Fig. 3H), but was not affected in the high gamma band (supplementary figure 8). Neither the amplitude nor the duration of gamma episodes was dependent on the intensity of optical stimulation. Indeed, basal forebrain optogenetic stimulation was adjusted to low (10 mW) or high (15 – 25 mW) laser power values. Only the mean frequency was marginally, yet significantly increased for slow gamma events (supplementary figure 8). Other parameters of gamma band episodes were not affected by optical stimulation of the basal forebrain (supplementary table 1, supplementary figure 8). Overall, these results support an active role for basal forebrain somatostatin cells in the regulation of cortical oscillatory activity, including slow waves and gamma band oscillations.

### Locomotor activity is triggered by basal forebrain disinhibition during resting states

Finally, we assessed the role of basal forebrain somatostatin cells in spontaneous behavioral patterns. For this, we tracked by video recording locomotor activity of freely-moving mice bilaterally implanted with optic fibers targeting the basal forebrain (supplementary figure 9, supplementary movies 1-3). Animals were placed in an open field and allowed to explore freely the environment. Once mice stopped exploration and became quiescent, we started optogenetic stimulation to the basal forebrain (Fig. 4A, B). Locomotor responses to laser stimulation of double transgenic NpHR+ animals were larger and faster than responses of control NpHR- animals (Fig. 4C). Locomotor responses were only different between NpHR+ and NpHR- animals when the initial state was quiescent (i.e.; not moving). Indeed, when animals were already moving, optogenetic inactivation of basal forebrain somatostatin cells produced no significant differences in locomotor displacements (supplementary figure 10). Furthermore, in order to discard nonspecific effects due to sensory detection of laser light, we repeated optogenetic stimulation in transgenic NpHR+ animals, but physically blocked the light path between ferrules (Fig. 4A). In doing so, we found that optogenetic stimulation was only effective in triggering movement when light was allowed to pass through the optic fiber to the basal brain (Fig. 4C). Under those conditions, responses were faster and larger. Thus, optogenetic inactivation of basal forebrain somatostatin neurons selectively elicits locomotor activity in quiescent animals, likely due to general cortical activation and increased arousal.

**Figure 4.**
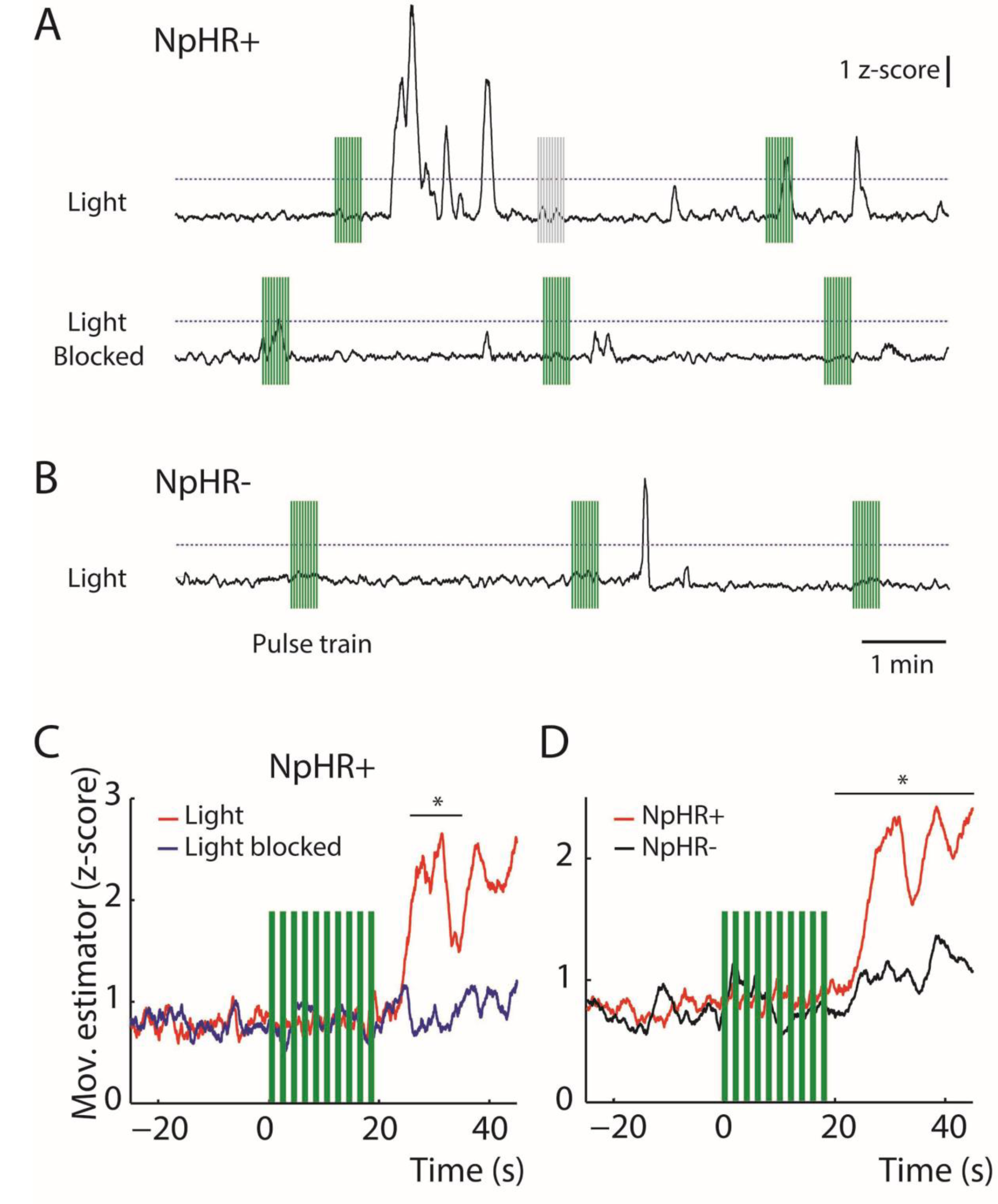
Locomotor activity following optogenetic inhibition of basal forebrain somatostatin neurons in resting mice. **A**, top panel: example of movement recorded over time in an NpHR+ animal upon laser stimulation of the basal forebrain. Bottom panel: movement is not induced when the light path is blocked between ferrules. **B**, example of a control NpHR animal stimulated by light delivered to the basal forebrain. Vertical bars depict laser train stimulation delivered when the animal is quiescent (green) or moving (grey) 40 s before stimulation. Dashed line indicates threshold (2 z-score) used to calculate movement amplitude. **C**, average response of NpHR+ animals (n = 3) to optogenetic inactivation of basal forebrain somatostatin cells with or without light path blockade. Response latency (from the end of stimulation) is significantly shorter when the light path is not blocked (seconds: 7.1 (range 0.3 – 40) vs 16.8 (range 0.3 – 40), p = 0.035, Wilcoxon rank-sum test). Response amplitude after the end of train stimulation is larger when the light path was not blocked (z-score: 2.96 (range 1.1 – 38.6) vs 2.28 (range 1.4 – 9.4), p = 0.004, Wilcoxon rank-sum test). **D**, average response of NpHR+ (n = 5 mice, 60 trials) and NpHR (n = 5, 49 trials) animals to optogenetic inactivation of basal forebrain somatostatin cells. Response latency (from the end of stimulation) is significantly shorter for NpRH+ animals (seconds: 11.8 (range 0.03 – 45) vs 45 (range 0.03 – 45), p = 0.0045, Wilcoxon rank-sum test). Response amplitude (after the end of train stimulation) is larger for NpHR+ animals (z-score: 2.92 (range 1.1 – 38.6) vs 2.27 (range 0.79 – 11.5), p = 0.0028, Wilcoxon rank-sum test). Note significant locomotor activation in NpHR animals- compared to the baseline, which likely results from nonspecific tissue heating during laser stimulation. Asterisks in C and D indicate statistical significance when the Wilcoxon rank-sum test was applied to compare both curves within a non-overlapping time window of 5 s. Pulse train; 10 1s pulses at 0.5 Hz, 15-20 mW.

## Discussion

We have shown that optogenetic inhibition of somatostatin cells in the basal forebrain is sufficient to locally modify the balance of synaptic activity and spiking patterns of some neuronal populations, therefore enhancing cortical activity and arousal. Such effect takes place rapidly (sub-second time scale) and likely involves the disinhibition of cholinergic and non-cholinergic pathways. Thus, somatostatin cells can exert a regulatory role on the synaptic output on the basal forebrain and indirectly control cortical processing and behavioral patterns.

### Optogenetic inactivation of somatostatin cells boosts the corticopetal synaptic output of the basal forebrain

Inhibition of neural spiking in somatostatin cells was rapidly followed by increased excitation of other neuronal populations, likely due to synaptic disinhibition (Ikeda and Wright, 1972). Given the fact that somatostatin cells provide functional inhibitory input to glutamatergic cells, cholinergic cells, and parvalbumin cells (Zaborszky and Duque, 2000, Xu et al., 2015), we believe that all these cell types might elevate their activity upon optogenetic inhibition of somatostatin cells. Interestingly, our data show that basal forebrain cells activated during laser stimulation had the highest baseline firing rates. Previous studies have shown that identified cholinergic cells exhibit low activity levels during slow wave sleep (Jones, 2005, Lee et al., 2005) and anesthesia-induced slow oscillations (Jones, 2005, Hassani et al., 2009). Accordingly, a significant proportion of our recorded units are likely to be cholinergic cells that were engaged during optical silencing of somatostatin cells. In addition, the local application of cholinergic receptor antagonists in the cortex confirmed that basal forebrain cholinergic cells were partially responsible for the laser induced effect of cortical activation. Taken together, this evidence suggests that inactivation of somatostatin cells is likely to recruit both cholinergic and non-cholinergic neurons in the basal forebrain.

### Enhancement and desynchronization of prefrontal cortex activity driven by the basal forebrain

Our results suggest that the synaptic drive provided by basal forebrain neurons disinhibited by optogenetic stimulation was powerful enough as to modify cortical activity patterns. The action of basal forebrain neurons on cortical dynamics is long known (Lin et al., 2015) and has been mostly attributed to cholinergic cells, despite the fact that they account for only a small fraction of basal forebrain neurons (Hedreen et al., 1984, Zaborszky et al., 2012). Our application of cholinergic receptor antagonists significantly reduced laser induced cortical activation, confirming that effects were partially mediated by cholinergic transmission. Enhanced neuronal discharge in the cortex during cholinergic activation has been reported in previous studies in vivo (Disney et al., 2007, Thiele et al., 2012, Pinto et al., 2013). Since we found little neuronal inhibition in the cortex during basal forebrain optical stimulation, our results suggest that the global effect of cholinergic transmission in the cortex might be shifting the balance of network activity to net excitation, with increased neuronal spiking. On the other hand, the decorrelation of cortical activity might be a general effect of cholinergic transmission in the cortex. Several observations support this idea. Indeed, during active whisking in mice, cholinergic fibers from the basal forebrain are robustly activated inducing transitions in cortical dynamics and brain state (Eggermann et al., 2014, Lin et al., 2015). Similarly, in the visual cortex, electrical stimulation of basal forebrain or optogenetic activation of cholinergic cells produces a marked decorrelation of neuronal spiking, which is associated with increased cognitive performance and enhanced sensory coding (Goard and Dan, 2009, Pinto et al., 2013). Finally, specific cholinergic lesions in the medial prefrontal cortex of monkeys have demonstrated the importance of basal forebrain cholinergic innervation for working memory (Croxson et al., 2011). Thus, we propose that enhanced neuronal discharge in the prefrontal cortex, characteristic of delay periods in working memory tasks (Fuster and Alexander, 1971, Kubota and Niki, 1971, Goldman-Rakic, 1990) is probably supported by fast cholinergic transmission. Moreover, we predict that neuronal spiking will exhibit reduced correlation within neighboring neurons selectively during delay periods. Future experiments will have to be designed to test these predictions.

### Alterations of network oscillations during cortical activation

Cortical slow oscillations occur during slow-wave sleep and deep anesthesia states, affecting large neuronal populations across the brain (Massimini et al., 2004). This rhythm is dichotomously organized into active and silent periods (Steriade et al., 1993b). During active periods (also called UP states (Sanchez-Vives and McCormick, 2000)) depolarized membrane potentials and elevated neuronal spiking predominate in cortical circuits; whereas during silent periods (also known as DOWN states (Sanchez-Vives and McCormick, 2000)) low synaptic activity and hyperpolarized membrane potential are largely synchronized (Steriade et al., 1993b, Sanchez-Vives and McCormick, 2000). Importantly, synaptic responsiveness (Timofeev et al., 1996) and sensory transmission (Azouz and Gray, 1999, Reig and Sanchez-Vives, 2007, Rigas and Castro-Alamancos, 2009) are differentially phase-modulated by the slow oscillation. However, results reported vary depending upon cortical areas studied and experimental protocols used. Accordingly, some studies suggest that active states might either enhance (Azouz and Gray, 1999, Reig and Sanchez-Vives, 2007) or decrease (Hasenstaub et al., 2007, Rigas and Castro-Alamancos, 2009) cortical responsiveness compared to silent states. Our results using optical stimulation of the basal forebrain suggest that responsiveness is enhanced in the medial prefrontal cortex during active states of the slow oscillation, possibly by exploiting neuronal depolarization and membrane fluctuations to amplify synaptic input (Destexhe et al., 2003, Reig et al., 2015). Similar neurophysiological mechanisms have been proposed for other cortical regions (Steriade, 2004, Munoz and Rudy, 2014). This is also supported by computational models that predict neuronal discharge exhibiting probabilistic behavior during active states, which modulates both synaptic gain and neuron transfer function (Ho and Destexhe, 2000, Destexhe and Contreras, 2006).

On the other hand, it has been shown that rhythmic neural activity in the gamma-frequency band in the medial prefrontal cortex is critical for several cognitive functions (Bosman et al., 2014); and alterations in their patterns have been associated with neuropsychiatric disorders, such as schizophrenia (Uhlhaas and Singer, 2010). Gamma oscillations in the cortex are locally generated by specific GABAergic cell populations (Cardin et al., 2009, Sohal et al., 2009). In addition, basal forebrain projection GABAergic cells can also contribute to enhance emergent gamma oscillations (Kim et al., 2015). These fast rhythms constitute a well-established marker of cortical activation and awake states (Steriade, 2004, 2006) that enhance neural circuit performance and information transfer between neurons (Sohal et al., 2009). We show here that optogenetic inhibition of basal forebrain somatostatin neurons suppresses slow waves and potentiates gamma oscillations in the cortex, which is consistent with the sleep-promoting role that has been proposed for basal forebrain somatostatin cells (Xu et al., 2015).

### Locomotor activity elicited by optogenetic stimulation of the basal forebrain during resting states

As a result of cortical activation, and given that the basal forebrain receives synaptic input from brainstem regions implicated in movement and arousal (Lee and Dan, 2012), we predicted that arousal and locomotion would also be increased during optogenetic inhibition of somatostatin cells. Moreover, a recent study has described a prominent projection from basal forebrain somatostatin cells directly to the ventral tegmental area and dorsal striatum (Do et al., 2016), reinforcing the idea that such neuronal population might be related with the regulation of locomotor activity and movement execution. Consistently, we found that bilateral inhibition of basal forebrain somatostatin cells elicited locomotor activity in quiescent animals. Since we did not monitor EEG or EMG activity, we cannot ascertain the stage of the sleep-wake cycle that animals were undergoing, yet it is possible that given the long periods of inactivity preceding optical stimulation (> 40 s), at least during some of the stimulation episodes, animals were sleeping. Interestingly, when animals were awake and active (i.e.; moving), optogenetic inactivation of basal forebrain somatostatin cells produced no significant differences in locomotor patterns. This suggests that the effects of basal forebrain activation triggered by somatostatin cells were strongly dependent on the ongoing behavioral state. Similar results have been previously reported for the effect of the basal forebrain on brain states. For example, in the visual cortex the effect of optogenetic activation of basal forebrain cholinergic neurons depends on the behavioral state immediately before the laser onset (Pinto et al., 2013). When animals are sitting still, cortical activity exhibits large-amplitude, low-frequency characteristic of quiet wakefulness (Crochet and Petersen, 2006). In such condition, basal forebrain activation causes a strong reduction of the low-frequency activity but no clear change at high frequencies. Instead, when animals are active (running), cortical activity shows less low-frequency activity, typical of active behavioral states (Crochet and Petersen, 2006), and basal forebrain activation causes a modest reduction of low-frequency power, but large increase at high frequencies (Pinto et al., 2013).

In summary, our results using optogenetic stimulation in the basal forebrain show that somatostatin cells are key elements in the regulation of local circuit activity, and can indirectly modulate cortical dynamics, producing increased neuronal spiking and decreased correlated discharge. Given their pivotal role in controlling basal forebrain synaptic output, somatostatin cells become a privileged target for the synaptic regulation of activity in the basal forebrain, which is at the crossroads of top-down and bottom-up regulatory pathways (Jones, 2005, Zaborszky et al., 2012). In this sense, the recently described long-range synaptic connectivity matrix of the basal forebrain (Do et al., 2016) is highly informative, as it provides rich anatomical information that will certainly help to understand the control mechanisms of basal forebrain activity. For example, the matrix of synaptic outputs provided by somatostatin cells is highly correlated with the synaptic inputs to all basal forebrain cell types (Do et al., 2016), suggesting that somatostatin neurons not only inhibit locally other cell types in the basal forebrain, but also suppress the exogenous input conveyed to those cell types; which is consistent with sleep-promoting role of somatostatin cells (Xu et al., 2015). Conversely, suppressed firing in somatostatin neurons in the basal forebrain will likely increase synaptic activity levels in the cortex, as we have found here, but also in other target areas.

## Methods

### Animals

All procedures involving experimental animals were performed in accordance to the U.S. Public Health Service’s Policy on Humane Care and Use of Laboratory Animals, reviewed and approved by university (CEBA) and national (CONICYT) bioethics committees. Experiments were carried out with 8- to 30-week-old mice (from either sex), in accordance with the Comité de Ética en Bienestar Animal (CEBA 13-014).

Three mice strains were used, C57Bl/6J (stock N° 000664), Ai39 (stock N° 014539, B6, 129S-Gt(ROSA)26Sor^tm39(CAG-HOP/EYFP)Hze^/J), and Sst-IRES-Cre (stock N° 013044, Sst^tm2.1(cre)Zjh^/J and stock N° 018973, B6N. Cg-Sst^tm2.1(cre)Zjh^/J). All transgenic lines were obtained from Jackson laboratories (www.jax.org). We used these strains as controls and refer to them as (Natronomonas pharaonis halorhodopsin) NpHR- animals throughout the text. Double transgenic animals were obtained from breeding Sst-IRES-Cre and Ai39 mice, so that they expressed functional NpHR+ exclusively in somatostatin cells. We refer to such animals as NpHR+ throughout the text. Mice were genotyped by PCR on ear biopsies using the primers: GGG CCA GGA GTT AAG GAA GA (Common), TCT GAA AGA CTT GCG TTT GG (Wild type Forward), TGG TTT GTC CAA ACT CAT CAA (Mutant Forward) for CRE Mice, and CTT TAA GCC TGC CCA GAA GA (Wild type Reverse), ATA TCC TGC TGG TGG AGT GG (Mutant Forward), GCC ACG ATA TCC AGG AAA GA (Mutant Reverse), TCC CAA AGT CGC TCT GAG (Wild type Forward) from Integrated DNA Technologies.

### In vivo electrophysiological recordings

Animals were induced with isoflurane, and then anesthetized with a dose of urethane (0.8 g/kg), and after 20 minutes a dose of ketamine (40 g/kg)/ xylazine (4 g/kg) to start the surgical procedures (Negron-Oyarzo et al., 2015). Throughout the experiment 1/12 of the initial dose of urethane was administered every 20-30 minutes. All drugs were administered intraperitoneally. Rectal temperature was monitored throughout the experiment and was kept at 36 °C with a heating pad. Glucosaline solution was injected subcutaneously every 2 hours.

In fully anesthetized mice, the scalp was cut and retracted to expose the skull. Mice were then implanted with a customized lightweight metal head holder and the head was held in a custome made metallic holder. Next, small craniotomies (~1 mm) were made with a dental drill above the basal forebrain (AP 0.38 mm and ML 1.5 mm from Bregma) (Franklin and Paxinos, 2007)and the prefrontal cortex (AP 2.5 mm and ML 0.35 mm from Bregma). The exposed dura was cut to expose the cortex giving access for implantation of the optic fiber and recording electrodes which were inserted at a depth of 4 mm and 1-2.2 mm, respectively. Neuronal activity in prefrontal cortex was recorded extracellularly with a 32 channel-four shank silicon probe (Buzsáki 32, Neuronexus) (mean resistance 1 MΩ) stained with DiI and inserted into the brain with a 30° angle towards the midline. Neuronal activity in basal forebrain was recorded by using a 32 channel-silicon probe (A1x32-Poly3-6mm-50-177, Neuronexus) stained with DiI and connected to an optic fibre (100 μm in diameter) attached to the shank (optrode). Electrical activity was recorded with a 32-channel Intan RHD 2132 amplifier board connected to an RHD2000 evaluation system (Intan Technologies). Single-unit activity and local field potential (LFP; sampling rate 20 kHz) were digitally filtered between 300 Hz – 5 kHz and 0.3 Hz – 2 kHz, respectively. Spike shape and amplitude were monitored during recording to ensure that the same cells were recorded.

To allow local drug injection a third craniotomy was made above mPFC (AP 2.0 mm and ML 0.87 mm from Bregma) and a 50 μm-tip pipette was inserted dorso-ventrally with a 30° angle towards the midline. For the blockade of cholinergic receptors in mPFC 200 nl of atropine (2 mM) and mecamylamine (2 mM) (1:1) (Sigma Aldrich) were microinjected at 1.4mm DV (IM-9B microinjector, Narishige), at minute 5 of the recording, while giving pulses of light on the basal forebrain and recording from mPFC.

### Surgery for chronic implantation

Mice were anaesthetized with isoflurane (5% induction and 1.5–2% maintenance) and placed on a stereotaxic frame (David Kopf Instruments). Temperature was kept at 37° throughout the procedure (1 – 2 hours) using a heating pad. The skin was incised to expose the skull and a craniotomy (~1 mm in diameter) was made with a dental drill above the basal forebrain bilaterally (AP +0.38 mm and ML +/-1.5 mm from Bregma) (Franklin and Paxinos, 2007). Two optic fibers (diameter 200 um, length 11 mm; Thorlabs) inserted and glued to ceramic ferrules (diameter 230 um, length 6.4 mm; Thorlabs) were descended through both craniotomies until reaching the basal forebrain and anchored to the skull using dental cement. After surgery, mice received a daily dose of enrofloxacin (10 mg/kg, Centrovet) for five days and supplementary analgesia with ketoprofen (5 mg/kg, Centrovet) for three days. Animals were allowed at least a week of recovery before behavioral tests.

### Optogenetic and somatosensory stimulation

Optogenetic stimulation of basal forebrain somatostatin neurons was achieved with a 200 μm optic fiber, (N. A. 0.37, Thorlabs) coupled to a green laser (532 nm) that provided a total light power of 0.1-60 mW at the fibre tip. An optrode was also used, which consisted of an optic fiber (100 μm, N. A. 0.22, Neuronexus) attached to an array of electrodes, so electrical recording and optical stimulation can be achieved simultaneously on the same site. Light stimuli consisted of 5 second light pulses and power at the tip of the fibre was set between 10 to 25 mW for 200 μm optic fibre and 4 to 6 mW for 100 μm optic fibre. A subset of experiments, both in NpHR+ and NpHR- animals, were performed with a 200 μm optic fibre and light power of 30 mW. At such intensity there was an evident increase in both the number of activated neurons and their discharge probability in NpHR- animals. Hence, the present study is based on experiments with light power up to 25 mW.

Somatosensory stimulation was applied by means of a tail pinch, with a solenoid (Takasago Electric) located on the tip of the rat’s tail and controlled by an Arduino UNO board (open-source microcontroller). Stimulus intensity (2 - 4 V output) and duration (typically 1 – 2 s) was manually adjusted to induce cortical activation, which was confirmed by the online visual inspection of the LFP frequency power content.

For chronically implanted animals, we randomly delivered a train (10 1-s square pulses at 1 Hz, 15 – 20 mW) every 2 – 3 min.

### Open field test

The testing arena (50 cm × 30 cm × 30 cm tall) was made of black painted acrylic, and illuminated by a 60 W bulb placed 150 cm above. The bilaterally implanted animal (see Surgery for details) was connected to optical fibers (200 um diameter, 1 m length) and placed on its cage to be habituated to the room for at least 15 min before testing. Next, the animal was placed in the testing arena for 10 to 20 minutes with no laser stimulation. This procedure was performed for three days to habituate the animal both to the room and arena. On the fourth day, the animal was placed in the testing arena for one hour and the stimulation protocol was then applied. The test was recorded using a digital video camera with a frame rate of 30 FPS. In some experiments, light transmission through the cannule was blocked with a small piece of aluminum foil placed between the ferrules.

### Behavioral event analysis

A custom MATLAB script was used to estimate animal movements. Briefly, the digital video was converted to a series of frames in RGB scale. For the whole video, consecutive frames on the green scale were used to detect the laser onset as well as to calculate the absolute value of the averaged difference between frames. Thus, any change in the image tracking could be quantified to estimate animal movements during the test. Finally, the movement estimator was z-scored to normalize different sessions and its absolute value was de-noised with a moving average (step = 50 bins). Laser effect was analyzed every time the animal was in a quiescent state before stimulation, i.e. movement estimator below 2 z-score for at least 40 s before the beginning of the laser train. The latency of the effect induced by laser was analyzed up to 40 s after the train stimulation offset (i.e. 60 s after the onset of laser train) and was calculated as the interval between 20 s and the time-point when the estimator exceeded 2 z-scores. Otherwise, latency was assumed to be 40 s.

### Histology and immunocytochemistry

At the end of recordings, mice were terminally anesthetized and intracardially perfused with saline followed by 20 min fixation with 4% paraformaldehyde. Brains were extracted and postfixed in paraformaldehyde for a minimum of 8 h before being transferred to PBS azide and sectioned coronally (60-70 μm slice thickness). Sections were further stained for Nissl substance. Location of shanks and optical fibre were determined in reference to standard brain atlas coordinates (Franklin and Paxinos, 2007) under a light transmission microscope.

For immunocytochemistry, non-recorded NpHR+ animals were terminally anesthetized and intracardially perfused with saline followed by 20 min fixation with 4% paraformaldehyde. Brains were extracted and sectioned coronally (60-70 μm slice thickness). Sections were rinsed three times for 10 min each with phosphate buffer, incubated in 1% horse serum supplemented with 0.3% Triton X-100 in phosphate buffer for 1 h, and then incubated in 1:2000 dilutions of the parvalbumin (PVG-2014, Swant) or ChAT (AB144P, Millipore) antibody for 24 h at 4°C, followed by a 1:1000 dilution of the secondary antibody for 3-6 hours at room temperature. Secondary antibodies were conjugated to Alexa Fluor 568 (Invitrogen); and cells were photographed with the appropriate filter cubes (Nikon; B-2E-C for EYFP, and G-2E/C for Alexa) with an epifluorescence microscope (Nikon Eclipse Ci). Antibody dilutions were performed in phosphate buffer with 1% horse serum and 0.3% Triton X-100. Sections were mounted on slides with mounting medium and photographed under epiluminescence microscopy. In NpHR+ animals, antibodies against somatostatin (ab108456, Abcam) failed under standard procedures. Thus, antigen retrieval was achieved by fixation with 3% paraformaldehyde solution, and heating sections at 80 °C for 10 minutes in citric acid (pH 6.0), prior to the procedure described above.

### Spike sorting

Semiautomatic clustering was performed by KlustaKwik, a custom program written in C++ (https://github.com/kwikteam/klustakwik2/). This method was applied over the 32 channels of the silicon probe, grouped in eight pseudo-tetrodes of four nearby channels. Spike clusters were considered single units if their auto-correlograms had a 2-ms refractory period and their cross-correlograms with other clusters did not have sharp peaks within 2 ms of 0 lag.

### Unit cross-correlation analysis

Neural activity in the cortex and basal forebrain was cross-correlated with the light pulse by applying the “sliding-sweeps” algorithm (Abeles and Gerstein, 1988). A time window of ± 15 s was defined with point 0 assigned to the light onset. The timestamps of the cortical and basal forebrain spikes within the time window were considered as a template and were represented by a vector of spikes relative to t = 0 s, with a time bin of 500 ms and normalized to the basal firing rate of the neurons. Thus, the central bin of the vector contained the ratio between the number of neural spikes elicited between ± 250 ms and the total number of spikes within the template. Next, the window was shifted to successive light pulses throughout the recording session, and an array of recurrences of templates was obtained. Both neural timestamps and start times of light pulses where shuffled by randomized exchange of the original inter-event intervals and the cross-correlation procedure was performed on the random sequence. The statistical significance of the observed repetition of spike sequences was assessed by comparing, bin to bin, the original sequence with the shuffled sequence. An original correlation sequence that presented a statistical distribution different from 100 permutations in at least three bins during the optical stimulation interval was considered as statistically significant, with p < 0.01 probability, instead of chance occurrence (see Statistics). If bins of the original correlation showed higher or lower values than the 100 permutations, neurons were classified as excited or inhibited by the light; respectively. Otherwise, neurons were identified as unaffected by optical stimulation.

### Spectral analysis

Time-frequency decomposition of LFP was performed with multi-taper Fourier analysis (Mitra and Pesaran, 1999) implemented in Chronux toolbox (http://www.chronux.org). LFP was downsampled to 500 Hz before decomposition. The same taper parameters described for the coherence analysis were used. To estimate gamma band power, spectra were normalized by 1/f (Mitra and Pesaran, 1999), in order to correct for the power law governing the distribution of EEG signals. To compute power and frequency of the slow/delta band oscillations, LFP was band-pass filtered with a two-way least squares FIR filter (0.5 - 2.0 Hz, eegfilt.m from EEGLAB toolbox; http://www.sccn.ucsd.edu/eeglab/); a Hilbert transform was applied and the mean value before (5 s) and during the light application (5 s) was calculated.

### Single-unit vs multi-unit coherence analysis

Single unit versus multi-unit coherence was determined using a previously described method (Pinto et al., 2013). Briefly, multi-unit activity was defined as the summed activity of all simultaneously recorded single units except the single unit used as reference for comparison. Spiking activity was then binned at 500 Hz and coherence for each single unit versus multi-unit pair was averaged for light ON (5 s) and light OFF (5 s before light onset) epochs. Coherence was computed using the multi-taper Fourier analysis (Mitra and Pesaran, 1999) as implemented in the Chronux toolbox (http://www.chronux.org). For each 5 s epoch, coherence was calculated using a time-bandwidth product of TW = 3 and 2TW-1 = 5 tapers, resulting in a half bandwidth W = 0.6 Hz. Wilcoxon signed rank test was applied to estimate the statistical significance of coherence results.

### Detection of cortical gamma band oscillations

A method developed to detect high-frequency oscillations in the hippocampus was modified to detect gamma band oscillations in the cortex (Logothetis et al., 2012). Briefly, cortical local field potential was downsampled (500 Hz) and band-pass filtered (20 – 80 Hz) using a zero phase shift non-causal finite impulse filter with 0.5 Hz roll-off. Next, the signal was rectified and low pass filtered at 20 Hz with a 4th order Butterworth filter. This procedure yields a smooth envelope of the filtered signal, which was then z-score normalized using the mean and SD of the whole signal. Epochs during which the normalized signal exceeded a 2 SD threshold were considered as events. The first point before threshold that reached 1 SD was considered the onset and the first one after threshold to reach 1 SD as the end of events. The difference between onset and end of events was used to estimate the gamma duration. We introduced a 150 ms-refractory window to prevent double detections. In order to precisely determine the mean frequency, amplitude, and duration of each event, we performed a spectral analysis using Morlet complex wavelets of seven cycles. Finally, a minimum duration criterion of 150 ms was used. The Matlab toolbox used is available online as LAN-toolbox (http://lantoolbox.wikispaces.com/).

### Statistics

Data sets were tested for normality using Kolmogorov-Smirnov test and then compared with the appropriate test (t-test or Wilcoxon two sided rank test). Statistical significance of data for protocols with factorial design (i.e., involving different contrasts and light on/off conditions) were assessed using two-way repeated-measures ANOVA followed by Bonferroni post-hoc comparison or Kruskal Wallis test followed by a by Mann-Whitney U contrasts. When necessary, significance analysis was estimated by applying the circ_corrcl.m in the CircStat toolbox of MATLAB (The Mathworks, Inc.) to calculate the p-value for correlation between one circular and one linear random variable. Hierarchical k-means cluster analysis was performed by using kmeans.m in in the Stats toolbox of MATLAB.

## Aknowledgments

We thank Drs. Mirian Schwalm, Daniel Rojas, and Hachi Manzur for reading and commenting on previous versions of this manuscript. This work was supported by the Comision Nacional de Investigacion Cientifica y Tecnologica (CONICYT) with grant Fondecyt regular 1141089 and grant CONICYT PIA ACT 1414.

## Supplementary Material

**Supplementary Figure 1.**
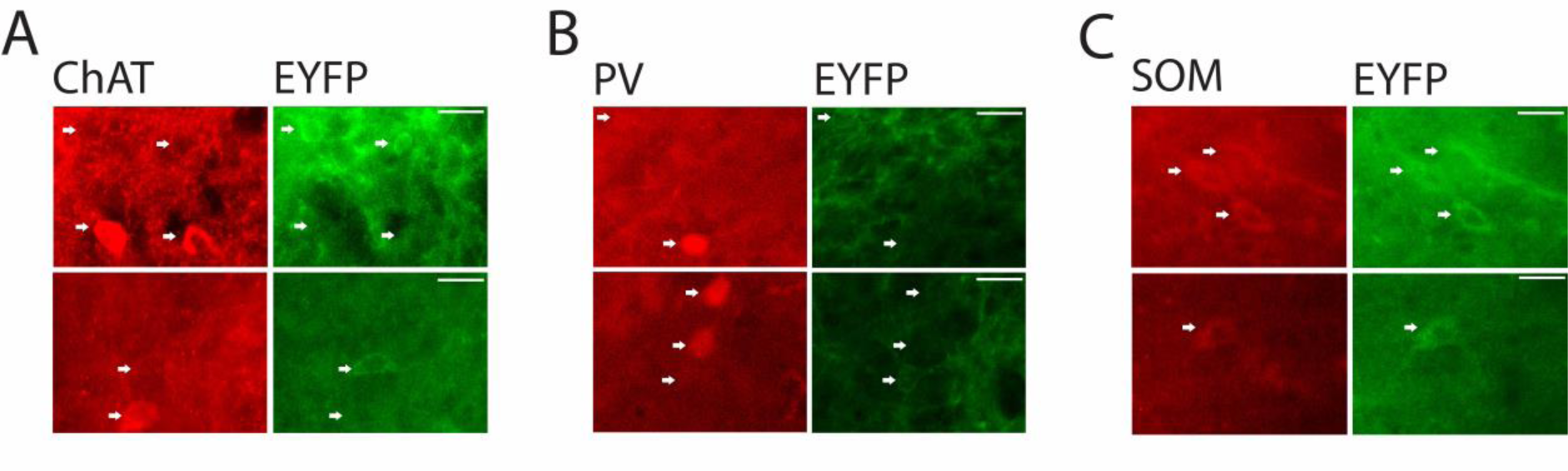
Halorhodopsin expression in the basal forebrain. Fluorescent micrographs showing the expression of choline acetyl transferase (ChAT, **A**), parvalbumin (PV, **B**), and somatostatin (SOM, **C**) in the basal forebrain of NpHR+ animals. EYFP depicts neurons expressing NpHR. Arrows depict neurons expressing one or both markers in each panel. Note no overlap between EYFP and ChAT (0%, n = 54 cells), little overlap between EYFP and PV (5%, n = 174 cells), and large overlap between EYFP and SOM (91%, n = 118 cells). Scale bar 20 um.

**Supplementary Figure 2.**
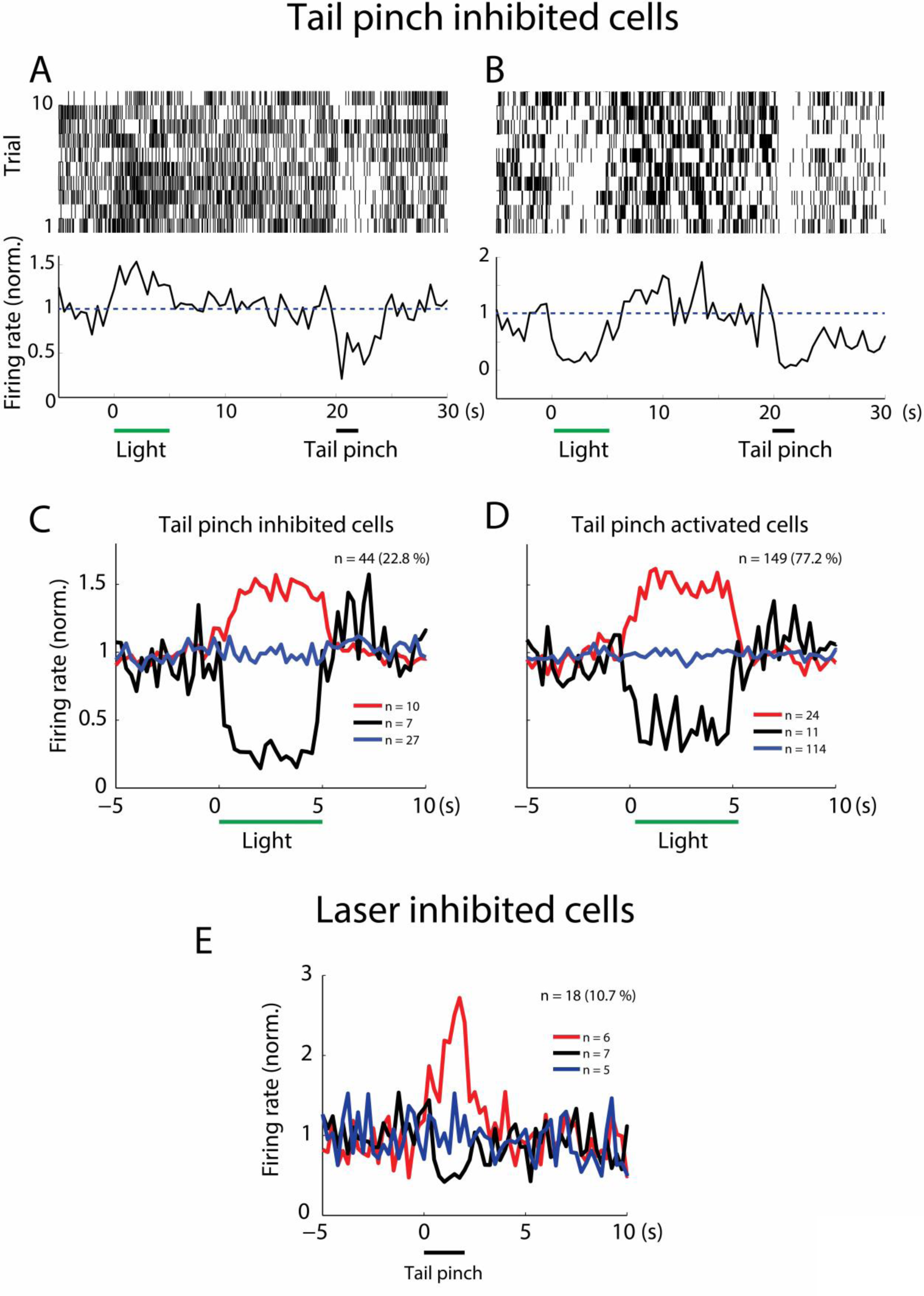
Local neuronal response patterns to combined optical and somatosensory stimulation in the basal forebrain. Examples of optogenetic excitation (A) or inhibition (B) of putative ChAT-negative cells, identified by their inhibition during somatosensory stimulation (tail pinch, (Hassani et al., 2009)). Top panels: raster plots; bottom panels: cumulative normalized discharge probability. Binsize: 500 ms. C, average normalized discharge probability for basal forebrain tail pinch-inhibited neurons (n = 44 cells, 4 animals). Cells were either excited (red line, n = 10), inhibited (black line, n = 7), or unaffected (blue line, n = 27) by light stimulation (horizontal green bar, 5 s, 15 – 25 mW). Binsize: 250 ms. D, average normalized discharge probability for tail pinch-excited neurons (n = 149 cells, 4 animals). Cells were either excited (red line, n = 24), inhibited (black line, n = 11), or unaffected (blue line, n = 114) by optical stimulation (horizontal black bar, 5 s, 15-25 mW). Binsize: 250 ms. E, average normalized discharge probability for basal forebrain somatostatin cells (n = 18 neurons, 2 animals). Cells were either excited (red line, n = 6), inhibited (black line, n = 7), or unaffected (blue line, n = 5) by somatosensory stimulation (horizontal black bar, 5 s, solenoid powered with 2 – 4 V).

**Supplementary Figure 3.**
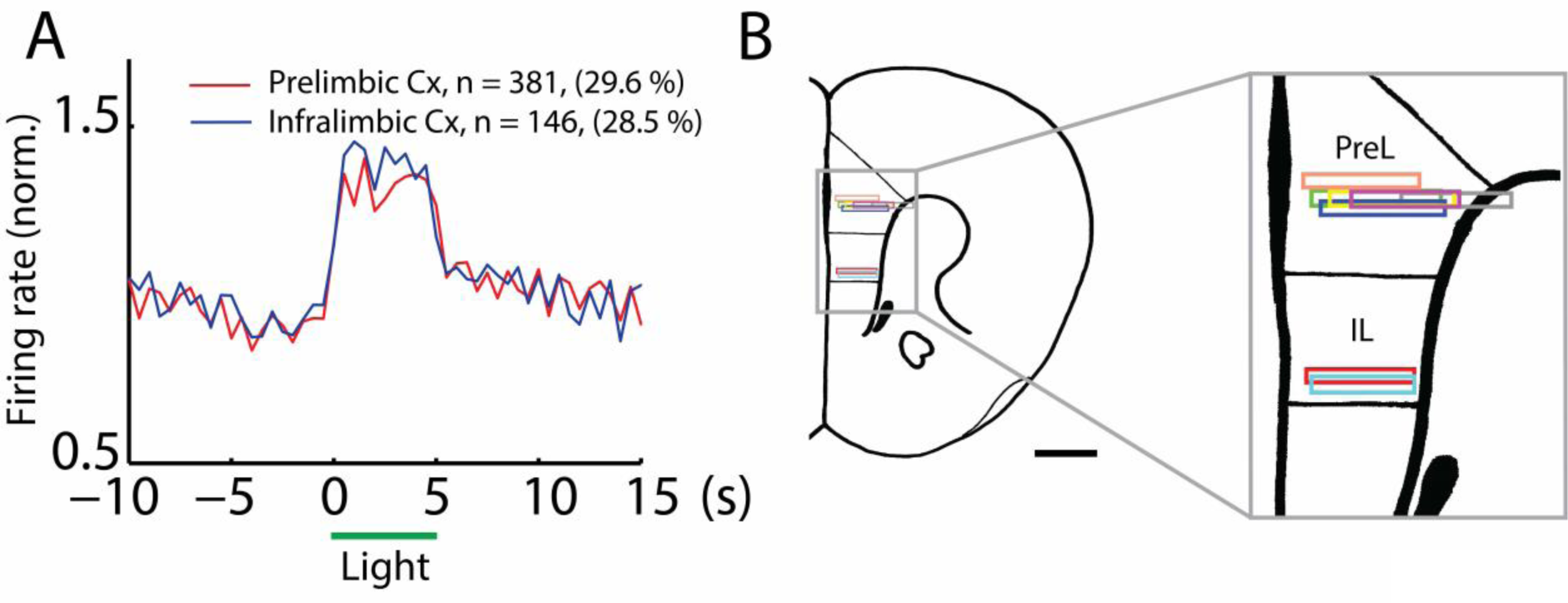
Effect of basal forebrain optogenetic stimulation on prelimbic and infralimbic regions of the prefrontal cortex. **A**, average normalized discharge probability of neurons excited by light on prelimbic (red line, n = 381 cells, 29.6%, 5 animals) or infralimbic (blue line, n = 146 cells, 28.5%, 2 animals) regions of the prefrontal cortex. Binsize: 500 ms. Horizontal green line depicts laser stimulation (5 s, 15 – 25 mW). Discharge probabilities were significantly different in prelimbic (1.36 ± 0.02) and infralimbic (1.31 ± 0.01) cortex (two-sample t-test, p = 0.0372). **B**, anatomical representation of recording locations (horizontal rectangles) in both cortical regions (n = 7 animals), based on plate 13 (Franklin and Paxinos, 2007). Scale bar: 1 mm.

**Supplementary Figure 4.**
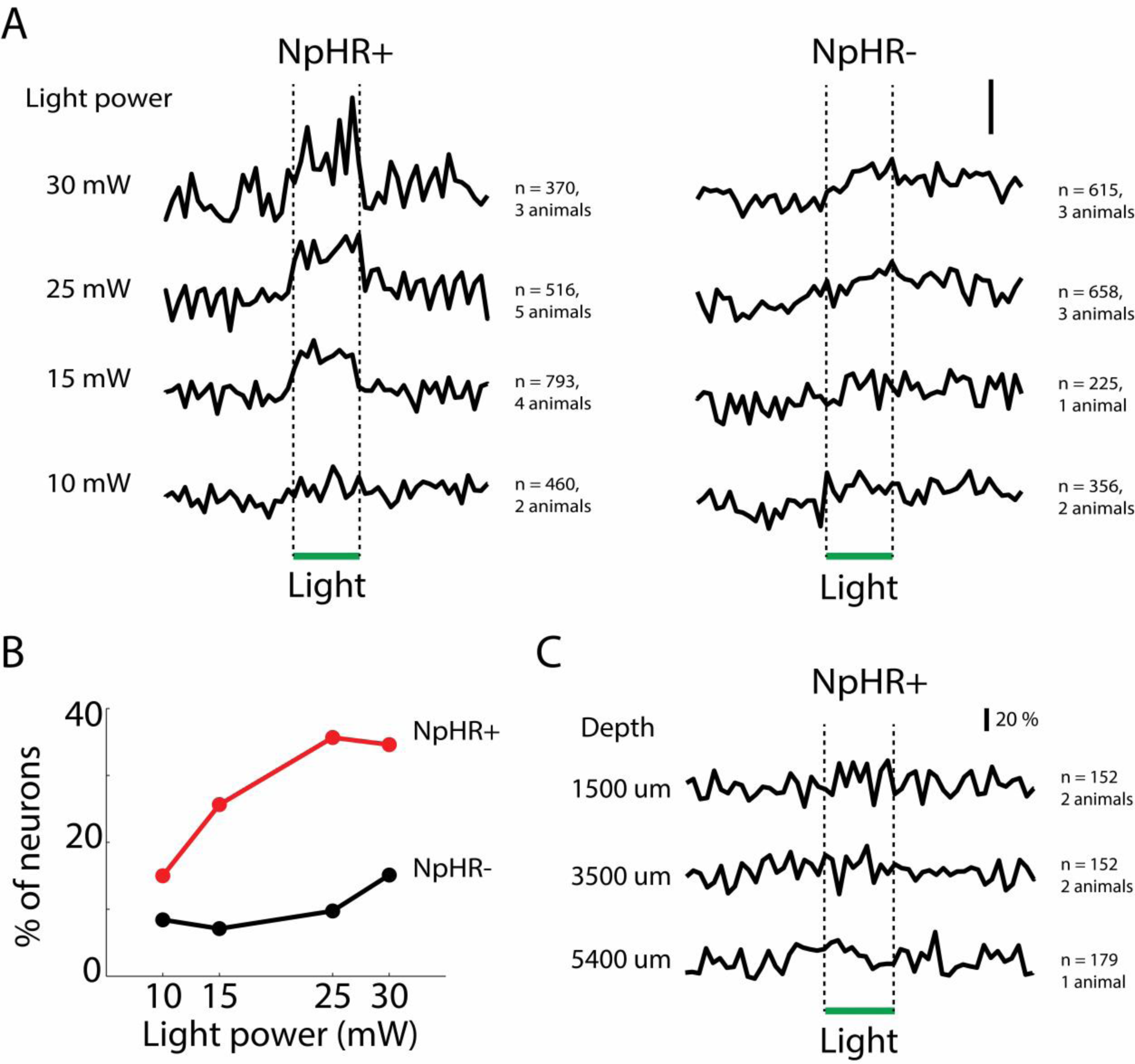
Neuronal spiking in the prefrontal cortex in response to optogenetic stimulation. **A**, average normalized discharge probability of cortical cells in NpHR+ (left column) and NpHR (right panel) animals. Horizontal bar indicates laser stimulation (5 s). Scale bar: 20% **B**, fraction of cortical neurons activated in relation to laser power on NpHR+ (red line) and NpHR- (black line) animals. Note that a minimum of ~10% of cortical neurons were activated in control animals (NpHR), with little effect on global spiking patterns. Nonspecific neuronal activation was likely due rise in temperature due to laser stimulation (Stujenske et al., 2015). **C**, Neuronal spiking in the prefrontal cortex in response to optogenetic stimulation in different anatomical locations in NpHR+ animals. Top trace, dorsal striatum (1500 um); middle trace, ventral striatum, (3500 um); bottom trace, olfactory tubercle, (5400 um). None of the responses was statistically significant (*p* > 0.05, Wilcoxon signed-rank test). Horizontal green bar depicts laser stimulation; 5 s, 15 – 25 mW

**Supplementary Figure 5.**
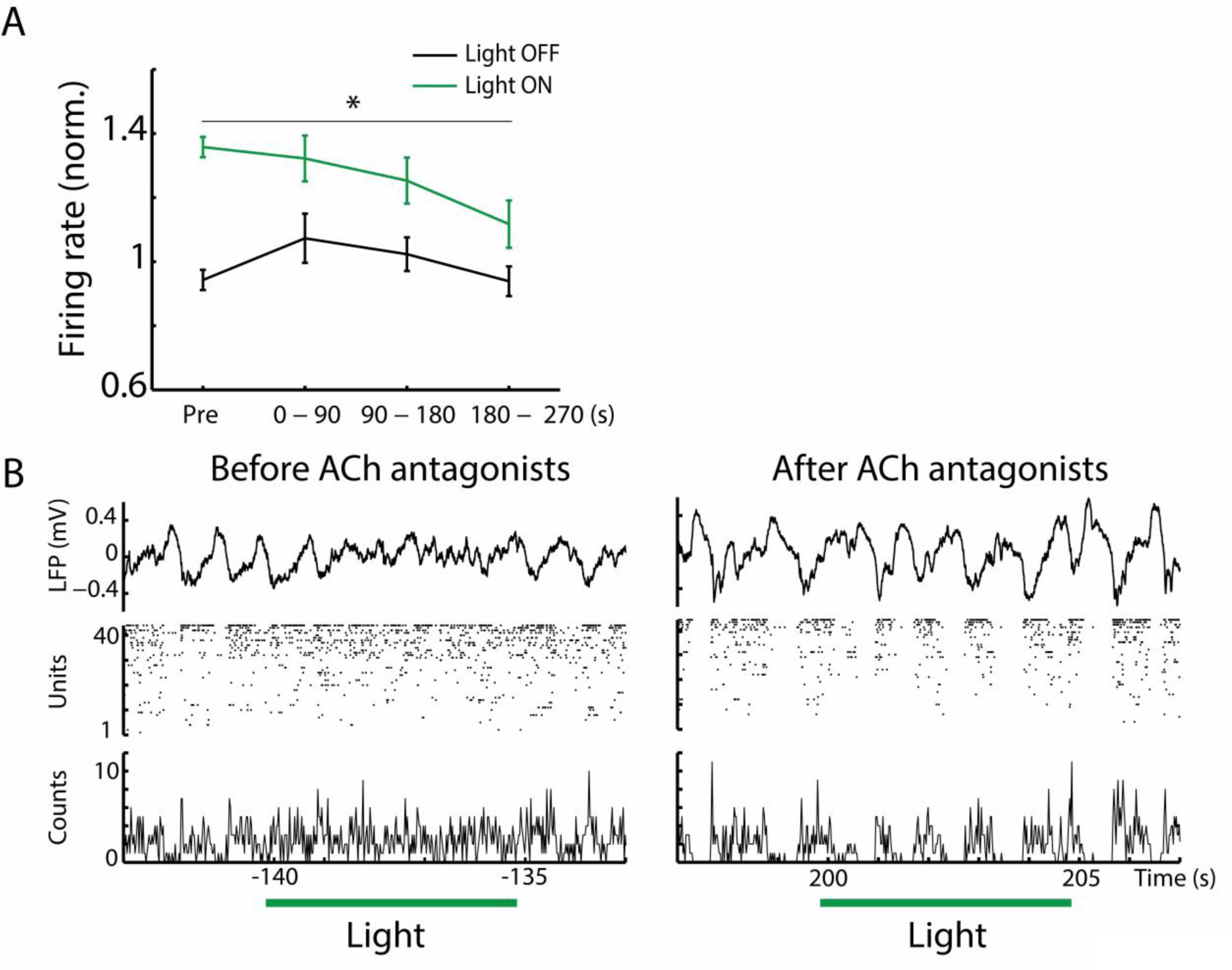
Role of cholinergic receptors in cortical spiking during optical inhibition of basal forebrain somatostatin cells. **A.** Average discharge probability at different time points in relation to drug application (time zero), in the presence (gray line) or absence (black line) of laser stimulation. **B.** example electrophysiological recording (NE84 reg12 chan1) showing response patterns before and after drug administration. Drug was injected at second zero (not shown). From top to bottom: LFP, raster plot for all cells, and multiunit histogram. Binsize: 2 ms. Horizontal green lines show laser stimulation (5 s, 25 mW).

**Supplementary Figure 6.**
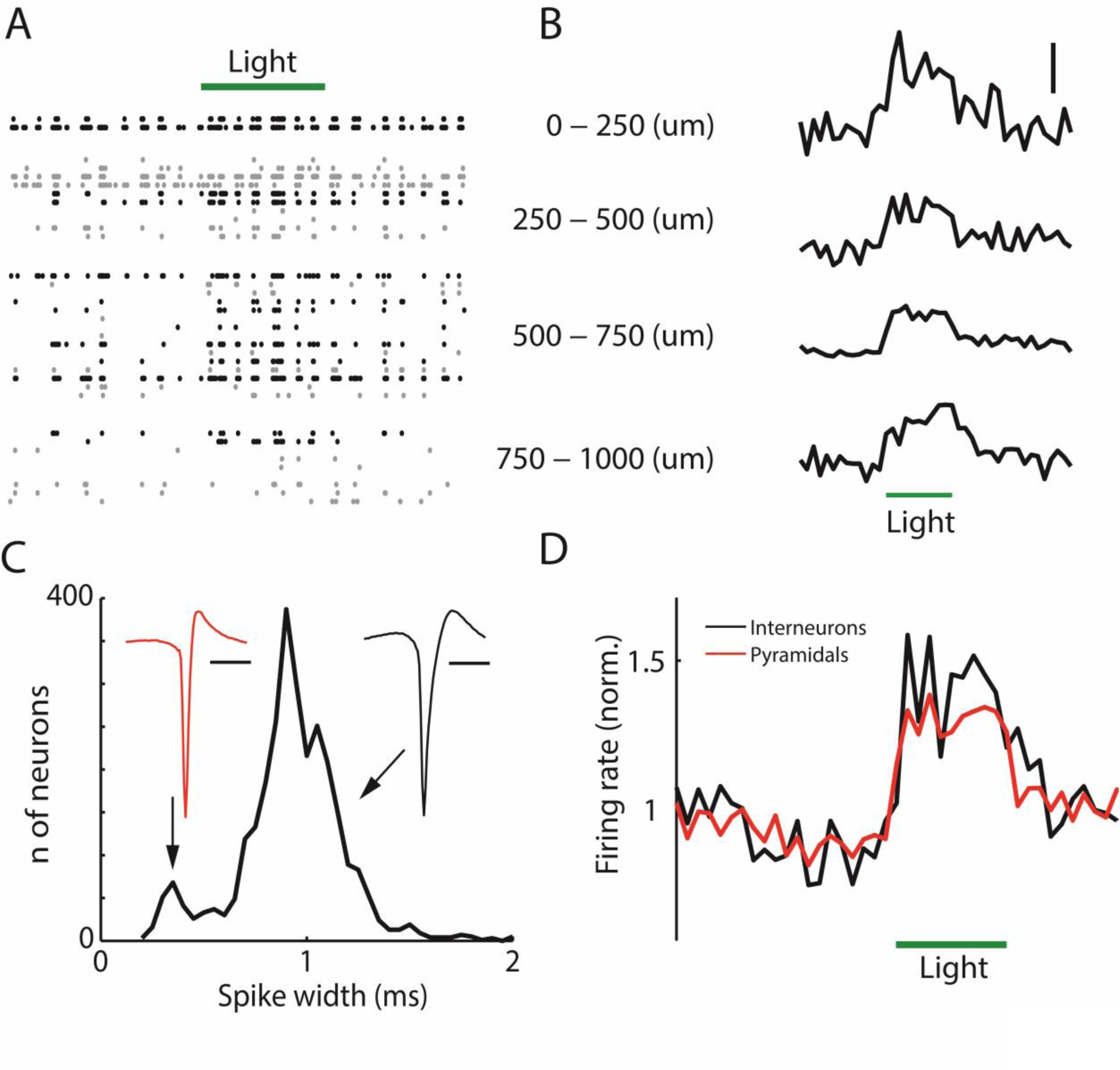
Cortical dynamics by layer and cell-type during optical inhibition of basal forebrain somatostatin cells. **A**, example raster plot for all units extracted from a cortical recording performed with a 4-shank 32-channel silicon probe. Note most recorded cells are located at intermediate depths. Black dots, laser-excited units; gray dots, unresponsive units. (NE83 reg14 chan1, 212 – 232 seconds) **B**, average normalized discharge probability of cortical neurons activated by basal forebrain optogenetic stimulation and recorded by depth. From top to bottom: n = 14 (25.5 %); n = 131 (30.8 %); n = 176 (28.7 %); n = 60 (31.1 %). Scale bar: 50 % **C**, histogram of spike durations (peak-to-trough) for all recorded cortical units (n = 2,895). Binsize: 50 us. Note bimodal distribution. Insets: average spike waveform for putative interneurons (red trace, width < 0.6 ms, n = 287, red waveform) and pyramidal cells (black trace, width > 0.6 ms, n = 2,588, black waveform). Scale bar: 1 ms. **D**, average discharge probability for interneurons (black line, n = 31) and pyramidal cells (red line, n = 350) excited by optical stimulation (\**p* < 0.05, Wilcoxon signed-rank test). Horizontal green lines show laser stimulation (5 s, 15 mW). Bin = 500 ms.

**Supplementary Figure 7.**
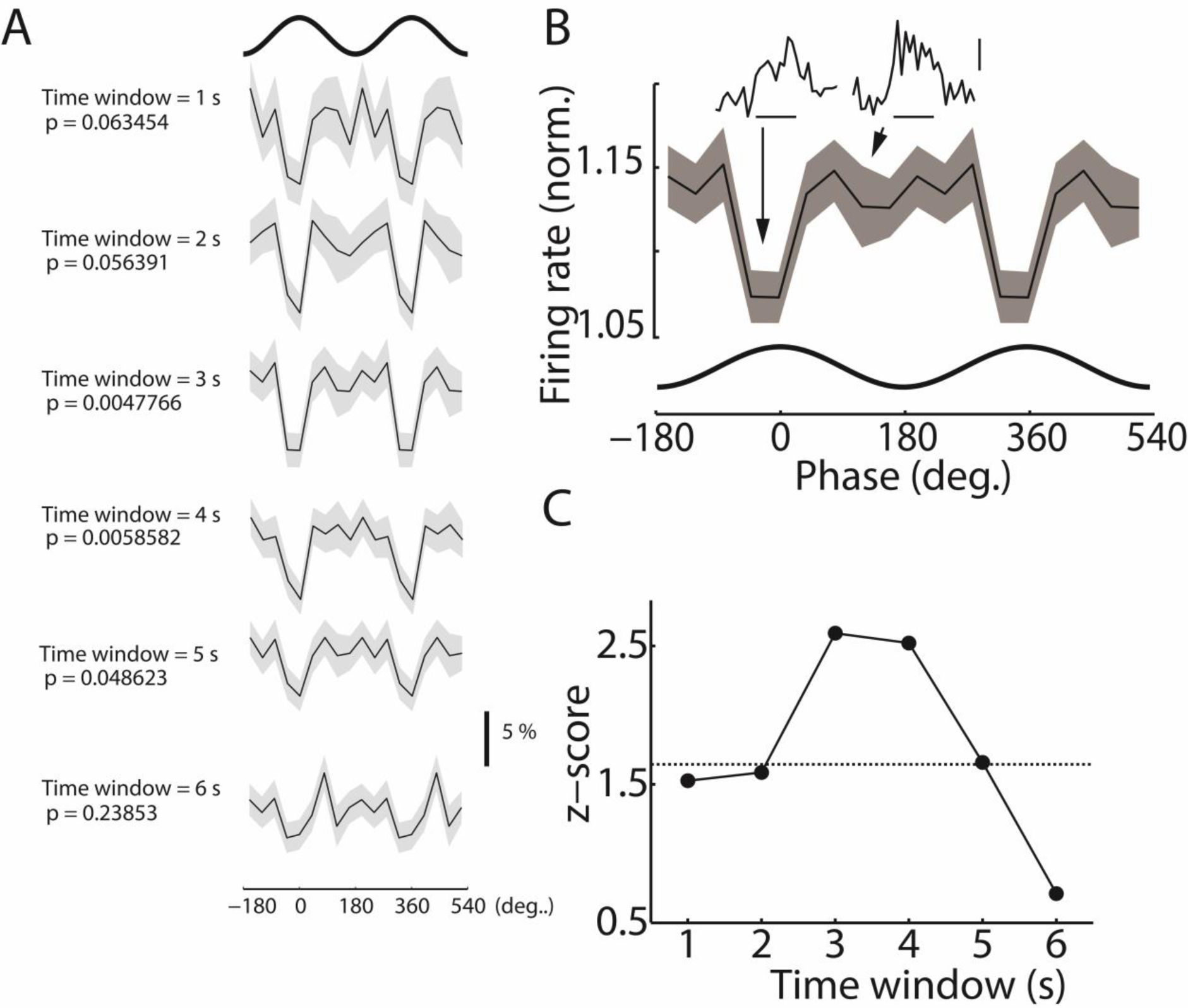
Phase-modulation of cortical activation during optical inhibition of basal forebrain somatostatin cells. **A**, phase histograms of average discharge probability induced by laser stimulation at different time windows. P-value is reported for different time windows starting at the light onset (circular-linear correlation). Solid line, mean; shaded area, s.e.m. Thick line on top of the figure represents two cycles of the slow oscillation. Scale bar: 5 %. The same data are repeated in two cycles for phase histograms to indicate oscillations. The peak of the extracellularly recorded oscillations in prefrontal cortex was at 0° and 360°. Bin size: 36°. **B**, phase histogram of average discharge probability induced by light stimulation (circular-linear correlation, *p* = 0.0047). Values were calculated during the first 3 seconds of the laser pulse. Black line shows mean and shaded area depicts s.e.m. Sinusoidal bottom line represents two cycles of the slow oscillation. The same data are repeated in two cycles for phase histograms to indicate oscillations. The peak of the extracellularly recorded oscillations in prefrontal cortex was at 0° and 360°. Bin size: 40°. Insets: average discharge probability profile for different phases of cortical slow oscillations. Left, light stimulation during cycle peaks (around 0 degrees). Right, light stimulation during cycle troughs (around 180 degrees). Note laser responses during peaks and troughs were significantly different (* *p* < 0.05, Wilcoxon signed-rank test). Scale bar: horizontal, laser stimulus (5 s); vertical, discharge probability (10 %). **C**, statistical significance z-score of the phase modulation discharge probability as a function of the integration time window (measured from laser pulse onset). Dashed line indicates statistical significance at *p* = 0.05. Note that differences in phase modulation were only significant during the second half of the laser pulse.

**Supplementary Figure 8.**
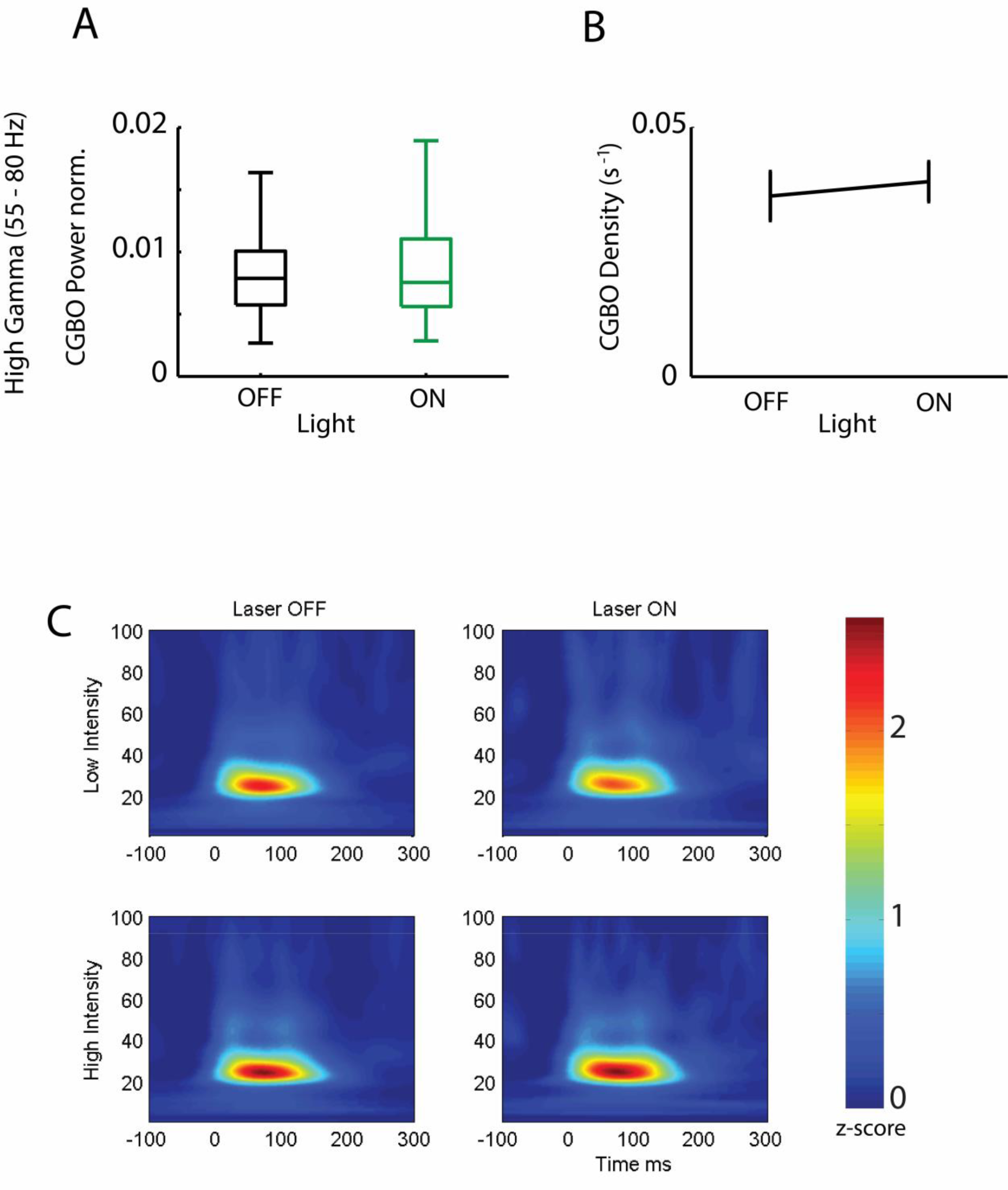
High gamma activity in the cortex during optical inhibition of basal forebrain somatostatin cells. **A**, box plots for the average power of high gamma band activity (55-80 Hz, *p* = 0.0520). **B**, plots (mean ± s.e.m) of density of high gamma band episodes (*p* = 0.17033). **C**, time resolved spectrogram of cortical activity for low cortical gamma band events. Basal forebrain optogenetic stimulation for low (10 mW) and high (15 – 25 mW) light intensities. Neither amplitude nor duration was affected by light intensity. Yet, the mean episode frequency was slightly increased (supplementary table 1).

**Supplementary Figure 9.**
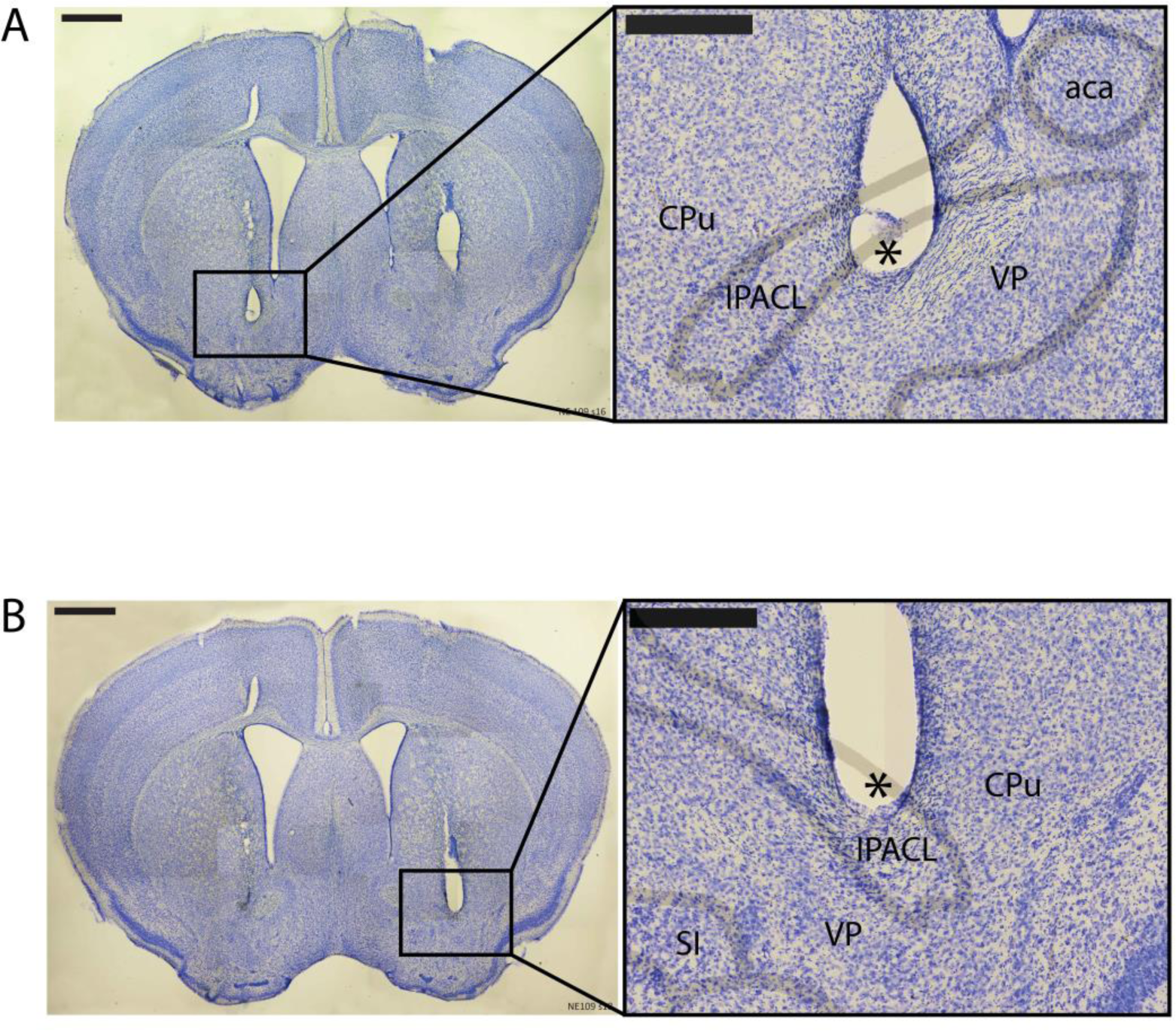
Photomontage of Nissl stained brain coronal sections from an example animal (NE109), showing the track and final tip position (insets) of bilateral optic fibers chronically implanted for optical stimulation experiments (Figure 4). Note both fibers reached the basal forebrain. **A**, section 18; **B**, section 16. Insets depict schematic borders of anatomical structures ( adapted from (Franklin and Paxinos, 2007)). Scale bars: thin line: 1 mm, thick line: 400 um. Asterisks indicate lesion. CPu, caudate putamen; IPACL: insterstitial nucleus of the posterior limb of the anterior commissure, lateral part; aca: anterior commissure, anterior; VP, ventral pallidum; SI: substantia innominata.

**Supplementary Figure 10.**
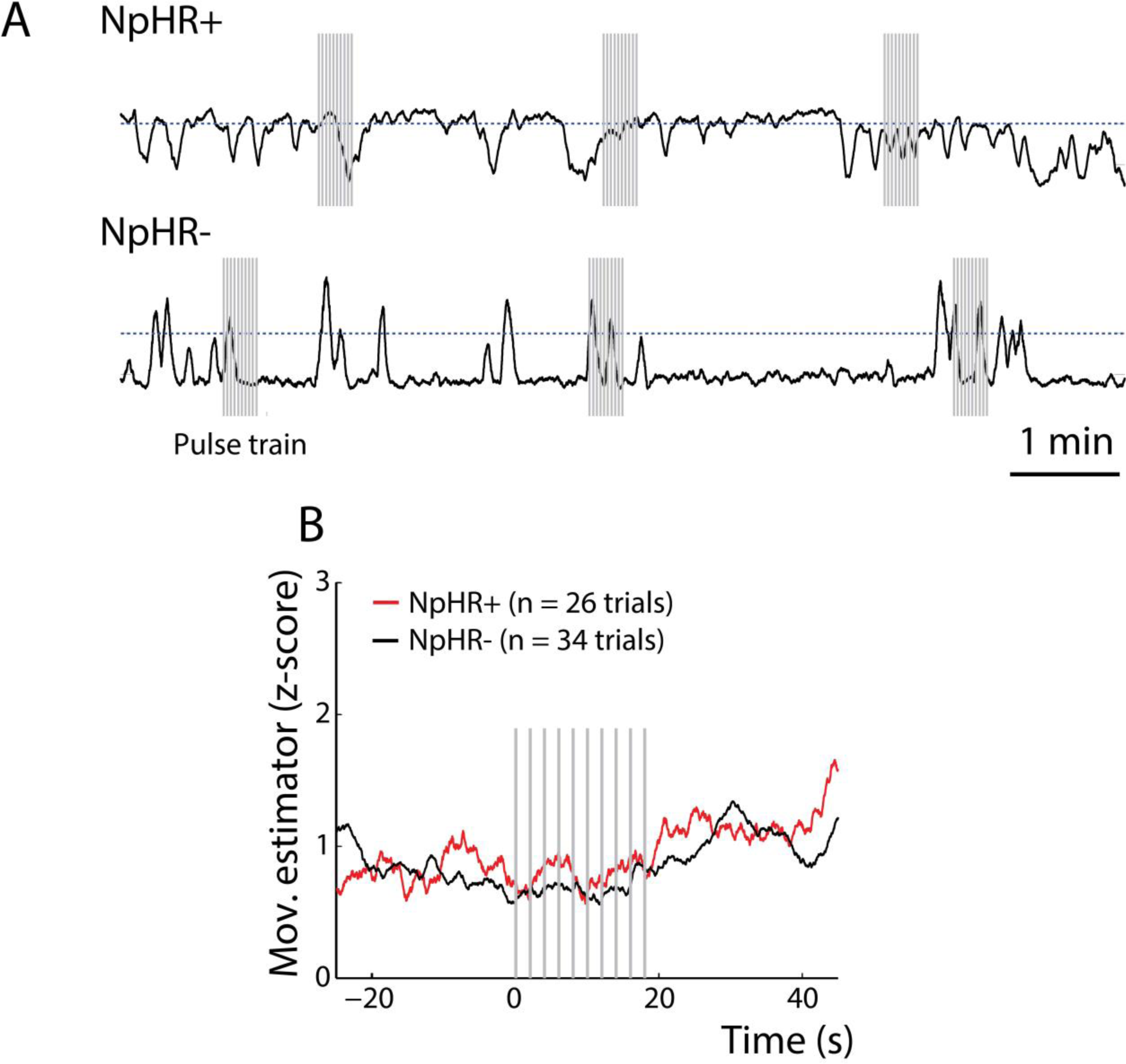
Locomotor activity upon optogenetic inhibition of basal forebrain somatostatin neurons in active mice. **A**, example of movement recorded over time in an NpHR+ (top panel) and NpHR- (bottom panel) animal upon laser stimulation of the basal forebrain. **B**, average responses of NpHR+ (n = 4) and NpHR- (n = 4) animals to optogenetic inactivation of basal forebrain somatostatin cells. Neither latency (p = 0.8) or amplitude (p = 0.22) of the response were significantly different between NpHR+ and NpHR- animals (Wilcoxon rank-sum test). Pulse train; 10 1s pulses at 0.5 Hz, 15-20 mW.

**Supplementary Table 1.** Parameters of gamma frequency events.

| Gamma oscillations |  |  | t-test |
| --- | --- | --- | --- |
| Low (20-40 Hz) | Light off | Light on | p - value |
| Duration (ms) | 156.44 ± 0.63 | 158.98 ± 1.42 | 0.0807 |
| Amplitude (z-score) | 1.44 ± 0.03 | 1.53 ± 0.05 | 0.0536 |
| Frequency (Hz) | 28.19 ± 0.10 | 28.65 ± 0.12 | 4.94E-04 |
| **High (55-80 Hz)** |  |  |  |
| **Duration (ms)** | 161.74 + 2.62 | 160.80 + 2.92 | 0.7739 |
| **Amplitude (z-score)** | 2.14 + 0.06 | 2.09 + 0.09 | 0.6298 |
| **Frequency (Hz)** | 68.19 + 0.42 | 68.96 + 0.65 | 0.2218 |

**Supplementary Movie 1.** (1337_ON_LD.avi) Example of behavioral effect induced by the optogenetic stimulation of the basal forebrain in a resting NpHR+ mouse in the open field.

**Supplementary Movie 2.** (1337_OFF_LD.avi) Example of behavioral effect induced by the optogenetic stimulation of the basal forebrain in a resting NpHR+ mouse in the open field with the light path blocked.

**Supplementary Movie 3.** (1289_CRE_LD.avi) Example of behavioral effect induced by the optogenetic stimulation of the basal forebrain in a resting NpHR- mouse in the open field.

